# Reward-driven changes in striatal pathway competition shape evidence evaluation in decision-making

**DOI:** 10.1101/418756

**Authors:** Kyle Dunovan, Catalina Vich, Matthew Clapp, Timothy Verstynen, Jonathan Rubin

**Author notes:** These authors contributed equally to this work (Kyle Dunovan, Catalina Vich). These authors contributed equally to this work (Timothy Verstynen, Jonathan Rubin). (TV). (JR).

## Abstract

Cortico-basal-ganglia-thalamic (CBGT) networks are critical for adaptive decision-making, yet how changes to circuit-level properties impact cognitive algorithms remains unclear. Here we explore how dopaminergic plasticity at corticostriatal synapses alters competition between striatal pathways, impacting the evidence accumulation process during decision-making. Spike-timing dependent plasticity simulations showed that dopaminergic feedback based on rewards modified the ratio of direct and indirect corticostriatal weights within opposing action channels. Using the learned weight ratios in a full spiking CBGT network model, we simulated neural dynamics and decision outcomes in a reward-driven decision task and fit them with a drift diffusion model. Fits revealed that the rate of evidence accumulation varied with inter-channel differences in direct pathway activity while boundary height varied with overall indirect pathway activity. This multi-level modeling approach demonstrates how complementary learning and decision computations can emerge from corticostriatal plasticity.

**Author summary:** Cognitive process models such as reinforcement learning (RL) and the drift diffusion model (DDM) have helped to elucidate the basic algorithms underlying error-corrective learning and the evaluation of accumulating decision evidence leading up to a choice. While these relatively abstract models help to guide experimental and theoretical probes into associated phenomena, they remain uninformative about the actual physical mechanics by which learning and decision algorithms are carried out in a neurobiological substrate during adaptive choice behavior. Here we present an “upwards mapping” approach to bridging neural and cognitive models of value-based decision-making, showing how dopaminergic feedback alters the network-level dynamics of cortico-basal-ganglia-thalamic (CBGT) pathways during learning to bias behavioral choice towards more rewarding actions. By mapping “up” the levels of analysis, this approach yields specific predictions about aspects of neuronal activity that map to the quantities appearing in the cognitive decision-making framework.

## 1 Introduction

The flexibility of mammalian behavior showcases the dynamic range over which neural circuits can be modified by experience and the robustness of the emergent cognitive algorithms that guide goal-directed actions. Decades of research in cognitive science has independently detailed the algorithms of decision-making (e.g., accumulation-to-bound models, [1]) and reinforcement learning (RL; [2, 3]), providing foundational insights into the computational principles of adaptive decision-making. In parallel, research in neuroscience has shown how the selection of actions, and the use of feedback to modify selection processes, both rely on a common neural substrate: cortico-basal ganglia-thalamic (CBGT) circuits [4–8].

Understanding how the cognitive algorithms for adaptive decision-making emerge from the circuit-level dynamics of CBGT pathways requires a careful mapping across levels of analysis [9], from circuits to algorithm (see also [10, 11]). Previous simulation studies have demonstrated how the specific circuit-level computations of CBGT pathways map onto sub-components of the multiple sequential probability ratio test (MSPRT; [5, 12]), a simple algorithm of information integration that selects single actions from a competing set of alternatives based on differences in input evidence [13, 14]. Allowing a simplified form of RL to modify corticostriatal synaptic weights results in an adaptive variant of the MSPRT that approximates the optimal solution to the action selection process based on both sensory signals and feedback learning [15, 16]. Previous attempts at multi-level modeling have largely adopted a “downwards mapping” approach, whereby the stepwise operations prescribed by computational or algorithmic models are intuitively mapped onto plausible neural substrates. Recently, Frank [17] proposed an alternative “upwards mapping” approach for bridging levels of analysis, where biologically detailed models are used to simulate behavior that can be fit to a particular cognitive algorithm. Rather than ascribing different neural components with explicit computational roles, this variant of multi-level modeling examines how cognitive mechanisms are influenced by changes in the functional dynamics or connectivity of those components. A key assumption of the upwards mapping approach is that variability in the configuration of CBGT pathways should drive systematic changes in specific sub-components of the decision process, expressed by the parameters of the drift diffusion model (DDM; [1]). Indeed, by fitting the DDM to synthetic choice and response time data generated by a rate-based CBGT network, Ratcliff and Frank [18] showed how variation in the height of the decision threshold tracked with changes in the strength of subthalamic nucleus (STN) activity. Thus, this example shows how simulations that map up the levels of analysis can be used to investigate the emergent changes in information processing that result from targeted modulation of the underlying neural circuitry.

Converging lines of evidence from human and non-human animal experiments have identified the striatum as playing a critical role in the process of accumulating evidence during decision-making [19], particularly in the context of value-based decision tasks [17]. In accumulation-to-bound models like the DDM, the drift rate parameter controls the speed at which evidence accumulates in favor of one choice over another (see [20]). It remains unclear how the dual-pathway organization of corticostriatal inputs contributes to the representation and comparison of evidence for conflicting actions. Recent theoretical models have proposed that decision evidence may be encoded as a dynamic competition between the direct and indirect pathways within a single CBGT action channel [7, 21–23]. In this scenario, the strength of evidence for a given action would be computed as the likelihood ratio of two hypotheses: “execute” (direct pathway) versus “suppress” (indirect pathway). Indeed, due to the opposing influence of dopamine (DA) on the sensitivity of direct and indirect MSNs to cortical input, DA-mediated RL could sculpt this competition to bias decisions towards the behaviorally optimal target [15]. In addition to this form of *within*-channel competition, competition *between* action channels may also contribute to the drift rate, such that greater differences between the activation of opposing direct pathways lead to faster evidence accumulation.

To see how the state of corticostriatal synaptic weights can impact decision process parameters, we adopted an upwards mapping approach to modeling adaptive choice behavior across neural and cognitive levels of analysis (Figure 1). Using a preliminary spike-timing dependent plasticity (STDP; [24, 25]) simulation, we modeled how phasic DA feedback signals [26] can modulate the relative balance of corticostriatal synapses, thereby promoting or deterring action selection. The effects of learning on the synaptic weights were incorporated into a spiking model of the full CBGT network meant to capture the known physiological properties and connectivity patterns of the constituent neurons in these circuits [27]. The performance (i.e., accuracy and response times) of the CBGT simulations were then fit using a hierarchical DDM [28]. This progression from synapses to networks to behavior allows us to show how variability in striatal pathway competition can manifest behaviorally by mapping how specific features of striatal activity that result from reward-driven changes in corticostriatal synaptic weights could underlie parameters of the fundamental cognitive algorithms of decision-making.

**Fig 1.**
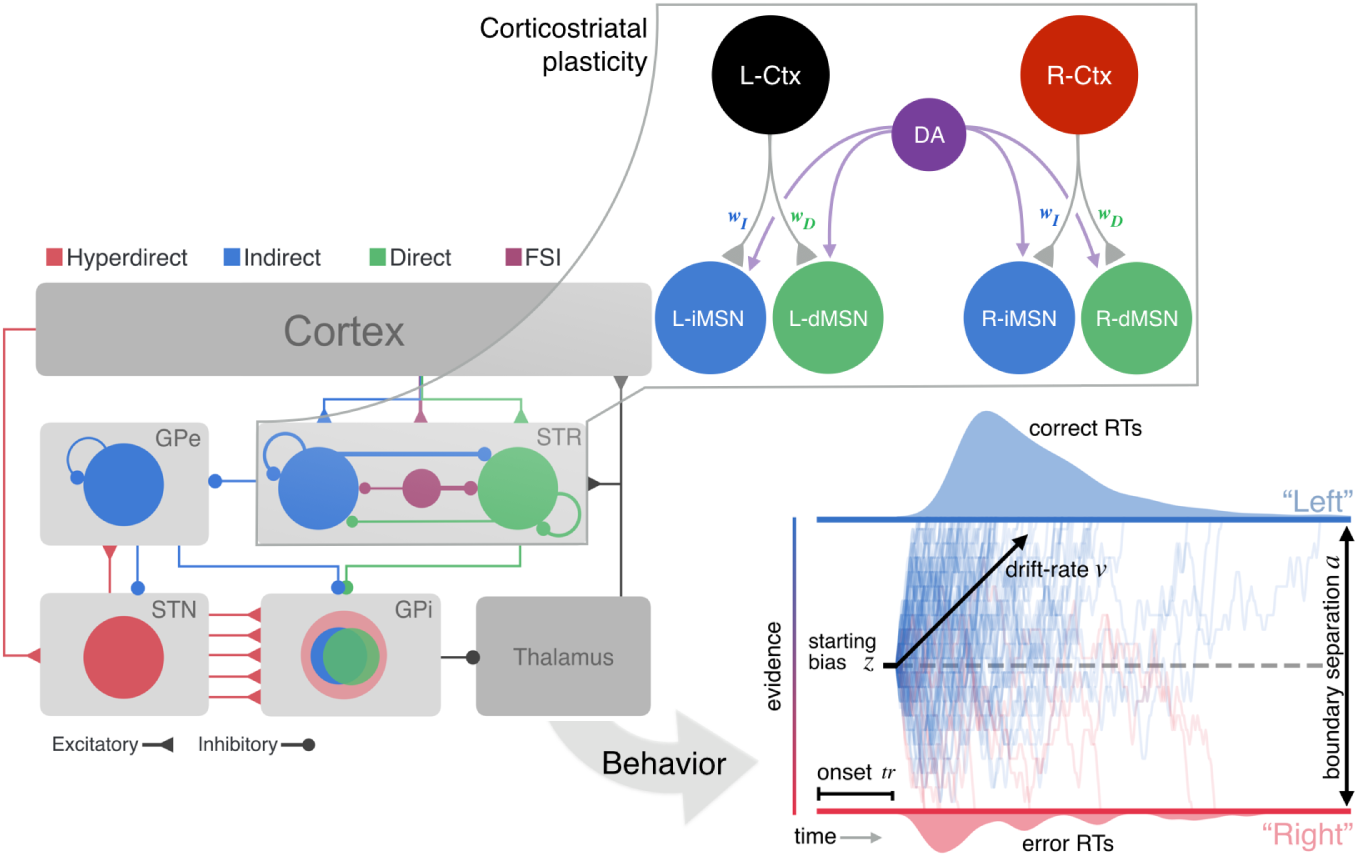
Multi-level modeling design. A STDP model of DA effects on Ctx-dMSN and Ctx-iMSN synapses is used to determine how phasic DA signals affect the balance of these synapses. A spiking model of the CBGT pathways simulates behavioral responses, under different conditions of Ctx-MSN efficacy based on the STDP simulations. The simulated behavioral responses from the full CBGT network model are then fit to a DDM of two-alternative choice behavior. Notation: *j − Ctx* - cortical population, *j − dMSN* - direct pathway striatal neurons, *j − iMSN* - indirect pathway striatal neurons (*j ∈ {L, R*}); DA - dopamine signal; STR - striatum; GPe - globus pallidus external segment; STN - subthalamic nucleus; GPi - globus pallidus internal segment; FSI - fast spiking interneuron; RT - reaction time; *v* - DDM drift rate; *a* - separation between boundaries in DDM; *z* - bias in starting height of DDM; *tr* - time after which evidence accumulation begins in DDM.

## 2 Results

### 2.1 Tuning corticostriatal synaptic weights

Our goal was to fit a DDM to behavioral data from our model CBGT network under different corticostriatal synaptic weight schemes. The first step was to find parameter settings for these synaptic weights, which are known sites of plasticity associated with dopaminergic signals about reward outcomes [26], that correspond to what would be observed after learning different levels of reward conflict in a two-alternative forced choice task. In this case, reward conflict is represented by the similarity in the reward probabilities associated with the two targets. In the absence of clear experimental consensus about how the strengths of direct and indirect pathway corticostriatal synapses vary across conditions, we followed the computational literature and performed a preliminary simulation of a reduced network with dopamine-related STDP in these weights [24, 25].

In the reduced network, one of two available actions, which we call left (*L*) and right (*R*), was selected by the spiking of model striatal medium spiny neurons (MSNs). These model MSNs were grouped into action channels receiving inputs from distinct cortical sources (Figure 1, left). Every time an action was selected, dopamine was released, after a short delay, at an intensity proportional to a reward prediction error based on comparison of the resulting reward to a value estimate from ongoing Q-learning. All neurons in the network experienced this non-targeted increase in dopamine, emulating striatal release of dopamine by substantia nigra pars compacta neurons, leading to plasticity of corticostriatal synapses (see Subsection 4.1 and [29]).

We first performed simulations in which a fixed reward level was associated with each action and confirmed that the reduced network agreed with several experimental benchmarks (see Supp. Figure 1): (a) firing rates in the direct pathway MSNs (dMSNs; firing rates *D_L_* and *D_R_*) associated with the more highly rewarded action increased, (b) firing rates of the indirect pathway MSNs (iMSNs; firing rates *I_L_* and *I_R_*) remained quite similar [30], and (c) both the dMSNs and the iMSNs associated with a selected action were active during the action selection process [31–33].

Given this confirmation of model performance, we next simulated a probabilistic reward task introduced in previous experimental studies on action selection with probabilistic rewards in human subjects [34]. For consistency with experiments, we always used *p_L_* + *p_R_* = 1, where *p_L_* and *p_R_* were the probabilities of delivery of a reward of size *r_i_* = 1 when actions *L* and *R* were performed, respectively. Moreover, as in the earlier work, we considered the three cases *p_L_* = 0.65 (high conflict), *p_L_* = 0.75 (medium conflict) and *p_L_* = 0.85 (low conflict). The simulation results show that corticostriatal synaptic weights onto the two dMSN populations clearly separated out over time (Figure 2). The separation emerged earlier and became more drastic as the conflict between the rewards associated with the two actions diminished, i.e., as reward probabilities became less similar. Interestingly, for relatively high conflict, corresponding to relatively low *p_L_*, the weights to both dMSN populations rose initially before those onto the less rewarded population eventually diminished (Figure 2A,B). This initial increase likely occurred because both actions yielded a reward of 1, leading to a significant dopamine increase, on at least some trials. The weights onto the two iMSN populations, on the other hand, remained much more similar, exhibiting just slight decreases over time (Figure 2A-C). The ratio of the dMSNs relative to the iMSNS, *w_D_/w_I_*, provided a single summary representation of the temporal evolution process, which highlighted the difference between channels within each reward scenario as well as the enhancement of this difference with less conflict between *p_L_* and *p_R_* (Figure 2D-F). We therefore extracted the corticostriatal synaptic weight ratios from the three probabilistic reward cases to use in our subsequent simulations of the CBGT network.

**Fig 2.**
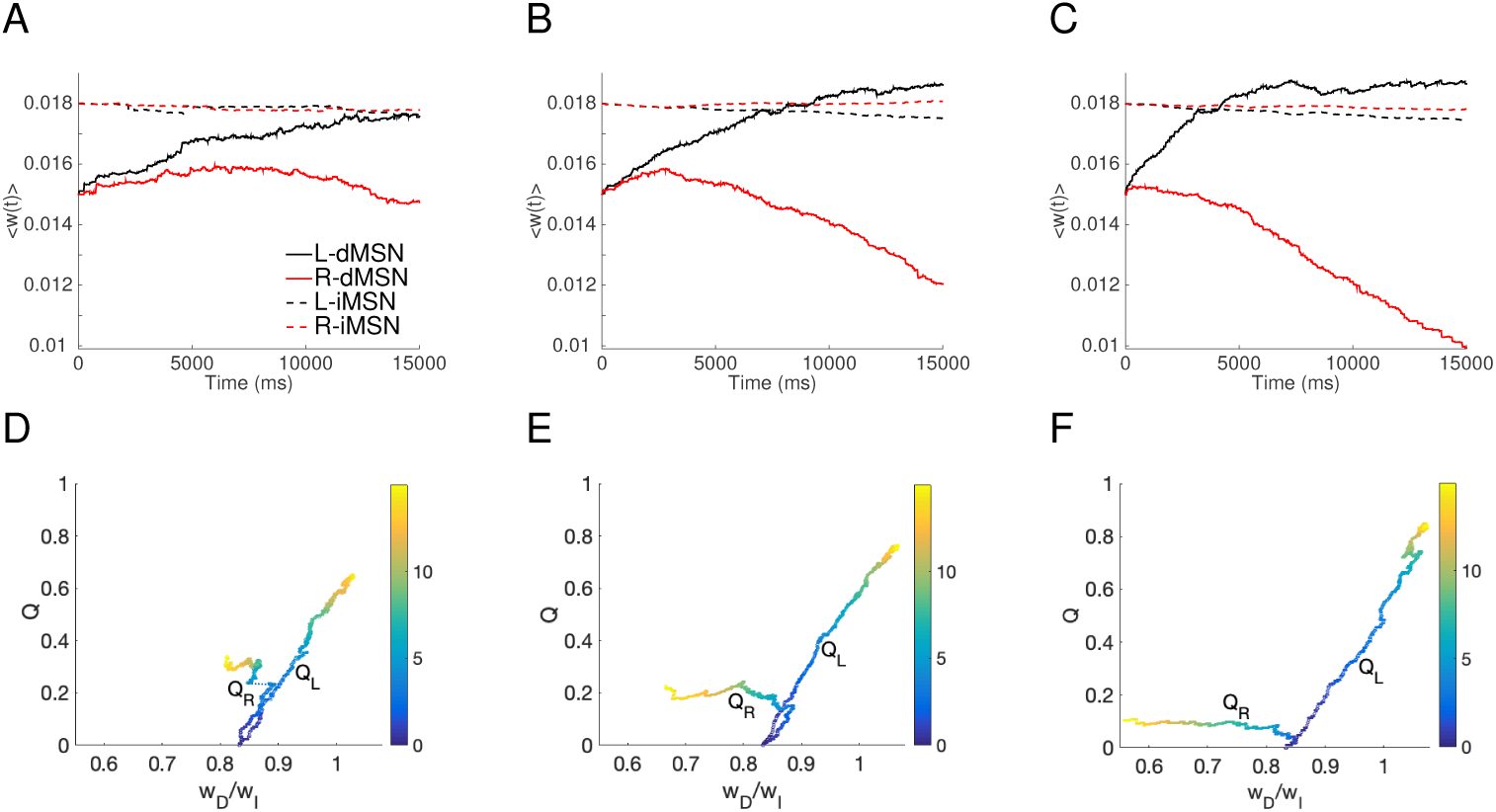
Corticostriatal synaptic weights with probabilistic reward feedback. First column: *p_L_* = 0.65; second column: *p_L_* = 0.75; third column: *p_L_* = 0.85 case. A, B, and C: Averaged weights over each of four specific populations of neurons, which are dMSN neurons selecting action *L* (solid black); dMSN neurons selecting action *R* (solid red); iMSN neurons countering action *L* (dashed black); iMSN neurons countering action *R* (dashed red). D, E, and F: Evolution of the estimates of the values for actions *L* (*Q_L_*) and *R* (*Q_R_*) estimated by Q-learning versus the ratio of the corticostriatal weights to those dMSN neurons that facilitate the action relative to the weights to those iMSN that interfere with the action. Both the weights and the ratios have been averaged over 8 different realizations. A small jump occurs in the *Q_R_* trace for *p_L_* = 0.65 and is joined by a dashed line; this comes from the time discretization and averaging.

### 2.2 CBGT dynamics and choice behavior

As illustrated by our STDP simulations, differences in rewards associated with different actions lead to differences in the ratios of corticostriatal synaptic weights, dMSN versus iMSNs, across action channels. Using weight ratios adapted from the STDP model, obtained by varying weights to dMSNs with fixed weights to iMSNs (Figure 2), we next performed simulations with a full spiking CBGT network (Figure 1) to study the effects of this corticostriatal imbalance on the emergent neural dynamics and choice behavior following feedback-dependent learning in the context of low, medium, and high probability reward schedules (2500 trials/condition; see Subsection 4.2.1 for details). In each simulation, cortical inputs featuring gradually increasing firing rates were supplied to both action channels, with identical statistical properties of inputs to both channels. These inputs led to evolving firing rates in nuclei throughout the basal ganglia, also partitioned into action channels, with an eventual action selection triggered by the thalamic firing rate in one channel reaching 30 *Hz* (Figure 3). We found that both dMSN and iMSN firing rates gradually increased in response to cortical inputs, and dMSN firing rates became higher in the channel for the selected action than for the other channel. Interestingly, iMSN firing rates also became higher in the selected channel, consistent with recent experiments (see [35], among others). Similar to the activity patterns observed in the striatum, higher firing rates were also observed in the selected channel’s STN and thalamic populations, whereas GPe and GPi firing rates were higher in the unselected channel (Figure 3).

**Fig 3.**
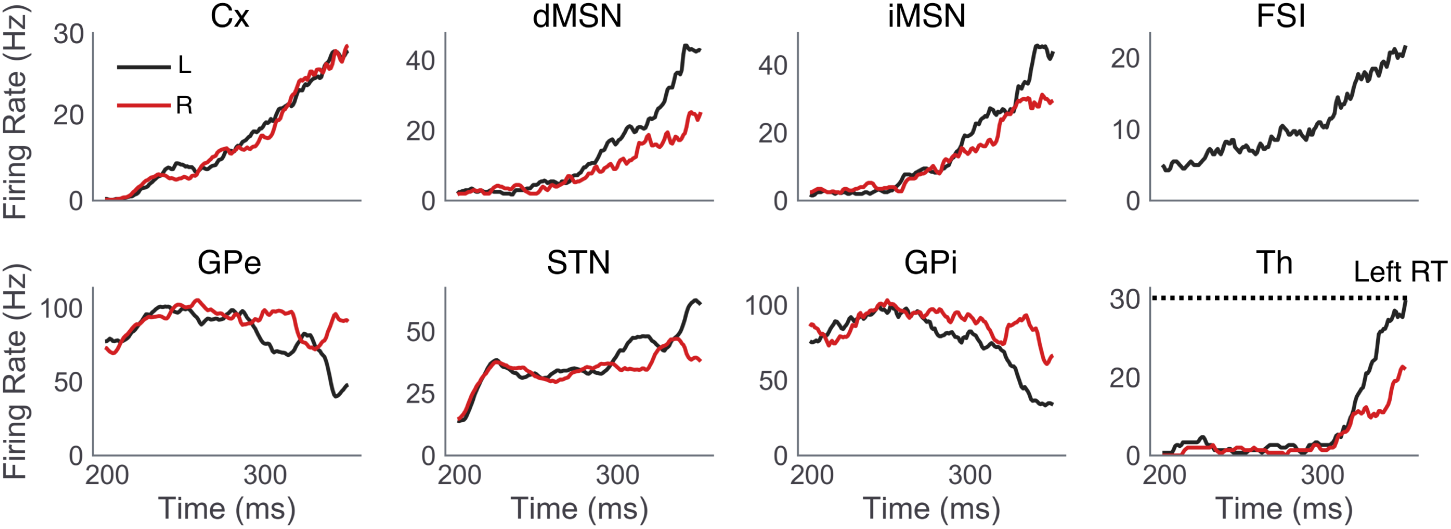
Single trial example of CBGT dynamics. Population firing rates of CBGT nuclei, computed as the average of individual unit firing rates within each nucleus in *L* (black) and *R* (red) action channels are shown for a single representative trial in the high reward probability condition. The selected action (*L*) and corresponding RT (324 *ms*) are determined by the first action channel to raise its thalamic firing rate to 30 *Hz*.

More generally across all weight ratio conditions, dMSNs and iMSNs exhibited a gradual ramping in population firing rates [19] that eventually saturated around the average RT in each condition (Figure 4A). To characterize the relevant dimensions of striatal activity that contributed to the network’s behavior, we extracted several summary measures of dMSN and iMSN activity, shown in Figure 4B-C. Summary measures of dMSN and iMSN activity in the *L* and *R* channels were calculated by estimating the area under the curve (AUC) of the population firing rate between the time of stimulus onset (200 *ms*) and the RT on each trial. Trialwise AUC estimates were then normalized between values of 0 and 1, including estimates from all trials in all conditions in the normalization, and normalized estimates were averaged over all trials. As expected, increasing the disparity of left and right Ctx-dMSN weights led to greater differences in direct pathway activation between the two channels (i.e., larger *D_L_ − D_R_*; Figure 4B). The increase in *D_L_ − D_R_* reflects a form of competition *between* action channels, where larger values indicate stronger dMSN activation in the optimal channel and/or a weakening of dMSN activity in the suboptimal channel. Similarly, increasing the weight of Ctx-dMSN connections caused a shift in the competition between dMSN and iMSN populations *within* the left action channel (i.e., an increase in *D_L_ −
I_L_*). Thus, manipulating the weight of Ctx-dMSN connections to match those predicted by the STDP model led to both between- and within-channel biases favoring firing of the direct pathway of the optimal action channel in proportion to its expected reward value.

**Fig 4.**
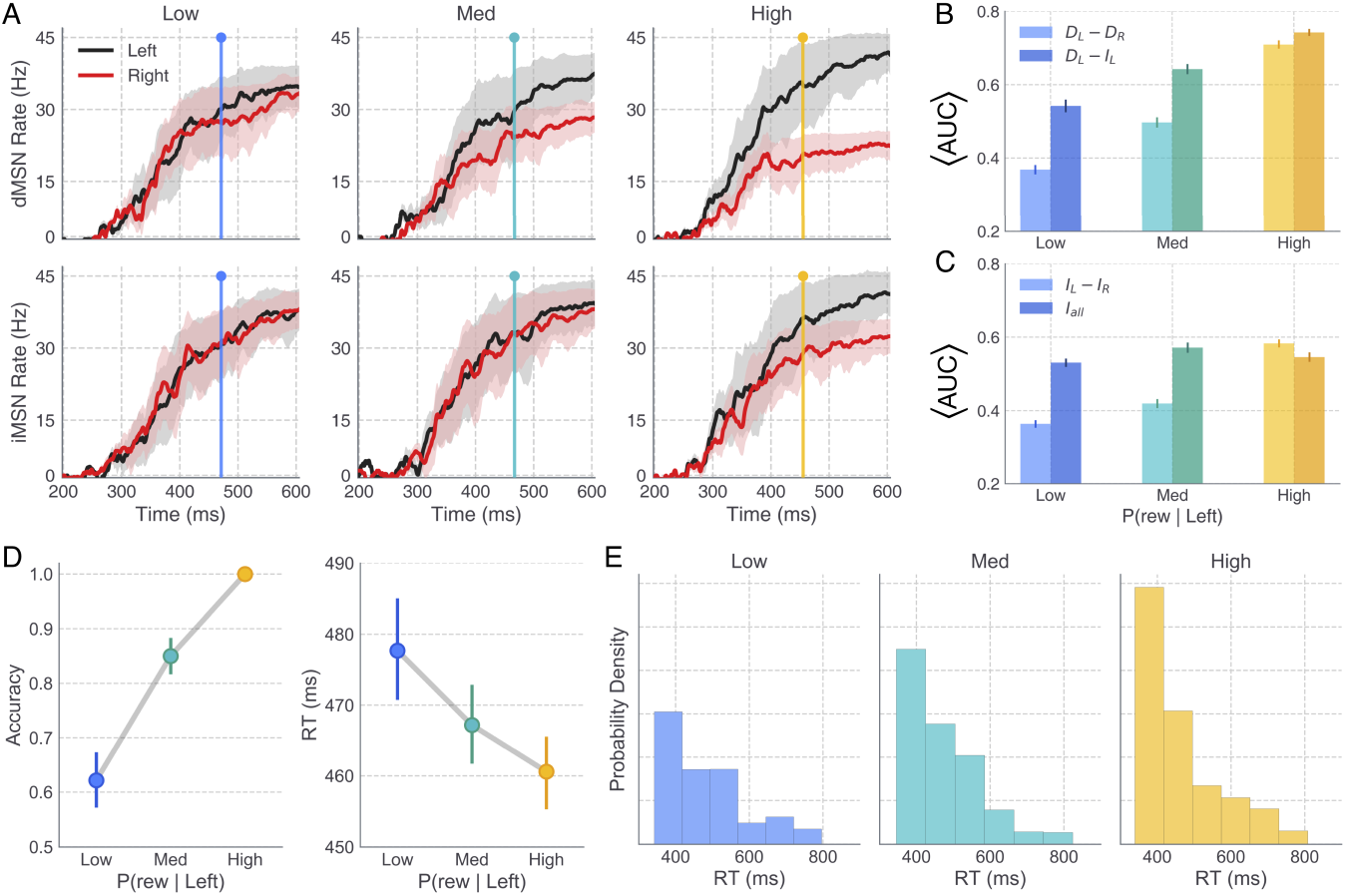
Striatal pathway dynamics and behavioral effects of reward probability in full CBGT network. A: Time courses show the average population firing rates for *L* (black) and *R* (red) dMSNs (top) and iMSNs (bottom) over the the trial window. Shaded areas reflect 95% CI. Colored vertical lines depict the average RT in the low (blue), medium (cyan), and high (yellow) reward conditions. B and C: Summary statistics of dMSN and iMSN population firing rates were extracted on each trial and later included as trialwise regressors on parameters of the DDM, allowing specific hypotheses to be tested about the mapping between neural and cognitive mechanisms. In B, lighter colored bars show the difference between dMSN firing rates in the *L* and *R* action channels whereas darker colored bars show the difference between dMSN and iMSN firing rates in the *L* action channel. Each was computed by averaging normalized values of trialwise estimates of the area under the appropriate firing rate curve (AUC); see main text for details. In C, lighter colored bars show the difference between iMSN firing rates in the *L* and *R* action channels and darker colored bars show the average iMSN firing rate (combined across left and right channels). Error bars show the bootstrapped 95% CI. D: Average accuracy (probability of choosing *L*) and RT (*L* choices only) of CBGT choices across levels of reward probability. E: RT distributions for correct choices across levels of reward probability; note that higher reward yields more correct trials. Error bars in B-D show the bootstrapped 95% CI.

Interestingly, although the weights of Ctx-iMSN connections were kept constant across conditions, iMSN populations showed reliable differences in activation between channels (Figure 4C). Similar to the observed effects on direct pathway activation, higher reward conditions were associated with progressively greater differences in the *L* and *R* indirect pathway firing rates (i.e., increased *I_L_ − I_R_*). At first glance, greater indirect pathway activation in more rewarded action channels seems to be at odds with the similarity of cortical synaptic weights to both indirect pathway channels that we obtained in the STDP model (Figure 2A) and also appears to be at odds with canonical theories of the roles of the direct and indirect pathways in RL and decision-making. This finding can be explained, however, by the presence of channel-specific excitatory inputs to the striatum from the thalamus in the CBGT network. That is, the strengthening and weakening of Ctx-dMSN weights in the *L* and *R* channels, respectively, translated into relatively greater downstream disinhibition of the thalamus in the *L* channel, which increased the excitatory feedback to *L*-dMSNs and *L*-iMSNs while reducing thalamostriatal feedback to *R*-MSNs in both pathways.

Finally, we examined the effects of reward probability on all iMSN firing rates, combined across action channels (*I_all_*; Figure 4C). Observed differences in *I_all_* across reward conditions were notably more subtle than those observed for other summary measures of striatal activity, with greatest activity in the medium reward condition, followed by the high and low reward conditions, respectively.

In addition to analyzing the effects of altered Ctx-dMSN connectivity strength on the functional dynamics of the CBGT network, we also studied how the decision-making behavior of the CBGT network was influenced by this manipulation. Consistent with previous studies of value-based decision-making in humans [36–40], we observed a positive effect of reward probability on both the frequency and speed of correct (i.e., leftward, associated with higher reward probability) choices (Figure 4D). Choice accuracy increased across low (*µ* = 64%, CI_95_ = [62, 65]), medium (*µ* = 85%, CI_95_ = [84, 86]), and high (*µ* = 100%, CI_95_ = [100, 100]) reward probabilities, *F* (2, 7497) = 702.38, *p <* 0.0001. Pairwise comparisons revealed that the increase in accuracy observed between low and medium conditions (*t*(4998) = 21.99, *p <* 0.0001), as well as that observed between medium and high conditions (*t*(4998) = 15.29, *p <* 0.0001), reached statistical significance. Along with the increase in accuracy across conditions, we observed a concurrent decrease in the mean RT of correct (*L*) choices in the low (*µ* = 477ms, CI_95_ = [472, 483]), medium (*µ* = 467ms, CI_95_ = [462, 471]), and high (*µ* = 460ms, CI_95_ = [456, 464]) reward probability conditions *F* (2, 6211) = 12.13, *p <* 0.0001. Notably, our manipulation of Ctx-dMSN weights across conditions manifested in stronger effects on accuracy (i.e., probability of choosing the more valuable action), with subtler effects on RT. Specifically, the decrease in RT observed between the low and medium conditions reached statistical significance (*t*(3712) = −2.9293, *p <* 0.0001); however, the RT decrease observed between the medium and high conditions did not (*t*(4624) = −2.0654, *p* = 0.13).

We also examined the distribution of RTs for *L* responses across reward conditions (Figure 4E). All conditions showed a rightward skew in the distribution of RTs, an empirical hallmark of simple choice behavior and a useful check of the suitability of accumulation-to-bound models like the DDM for modeling behavioral data sets. Moreover, the degree of positive skew in the RT distributions (i.e., greater probability mass at lower than at higher values) for *L* responses became more pronounced with increasing reward probability, suggesting that the observed decrease in the mean RT at higher levels of reward was driven by a change in the shape of the distribution, and not, for instance, a temporal shift in its location. While the skewed shape of the RT distributions produced by the CBGT network are qualitatively consistent with those typically observed in human choice experiments, it is worth noting that the network-simulated RTs appeared to follow an exponential distribution, whereas empirical RTs often exhibit a mode closer to the center of mass.

### 2.3 CBGT-DDM mapping

We performed fits of a normative DDM to the CBGT network’s decision-making performance (i.e., accuracy and RT) to understand the effects of corticostriatal plasticity on emergent changes in decision behavior. This process was implemented in three stages. First, we compared models in which only one free DDM parameter was allowed to vary across levels of reward probability (single parameter DDMs). Next, a second round of fits was performed in which a second free DDM parameter was included in the best-fitting single parameter model identified in the previous stage (dual parameter DDMs). Finally, the two best-fitting dual parameter models were submitted to a third and final round of fits with the inclusion of trialwise measures of striatal activity (see Figure 4B-C) as regressors on designated parameters of the DDM.

All models were evaluated according to their relative improvement in performance compared to a null model in which all parameters were fixed across conditions. To identify which single parameter of the DDM best captured the behavioral effects of alterations in reward probability as represented by Ctx-dMSN connectivity strength, we compared the deviance information criterion (DIC) of DDM models in which either the boundary height (*a*), the onset delay (*tr*), the drift rate (*v*), or the starting-point bias (*z*) was allowed to vary across conditions. Figure 5A shows the difference between the DIC score of each model (DIC_*M*_) and that of the null model (ΔDIC = DIC_*M*_ − DIC_*null*_), with lower values indicating a better fit to the data (see Table 1 for additional fit statistics). Conventionally, a DIC difference (ΔDIC) of magnitude 10 or more is regarded as strong evidence in favor of the model with the lower DIC value [41]. Compared to the null model as well as alternative single parameter models, allowing the drift rate *v* to vary across conditions afforded a significantly better fit to the data (ΔDIC = −960.79). Examination of posterior distributions of *v* in the best-fitting single parameter model revealed a significant increase in *v* with successively higher levels of reward probability (*v_Low_* = 0.35; *v_Med_* = 1.61; *v_High_* = 2.71), capturing the observed increase in speed and accuracy across conditions by increasing the rate of evidence accumulation toward the upper (*L*) decision threshold.

**Table 1.** Single- and dual-parameter DDM goodness-of-fit statistics. DIC is a complexity-penalized measure of model fit, DIC = D(*θ*) + *pD*, where D(*θ*) is the deviance of model fit under the optimized parameter set *θ* and *pD* is the effective number of parameters. ΔDIC is the difference between each model’s DIC and that of the null model for which all parameters are fixed across conditions. Asterisks denote models providing best fits within the single-parameter group (*) and across both groups (**).

| | DIC | $\Delta DIC$ |
| --- | --- | --- |
| Null | -9144.21 | 0.0 |
| Bound ($a$) | -9177.03 | -32.83 |
| Onset ($tr$) | -9175.54 | -31.34 |
| Drift ($v$)* | -10105.0 | -960.79 |
| Bias ($z$) | -9447.5 | -303.29 |
| $v, a$** | -10318.27 | -1174.07 |
| $v, tr$ | -10113.38 | -969.17 |
| $v, z$ | -10134.22 | -990.02 |

**Fig 5.**
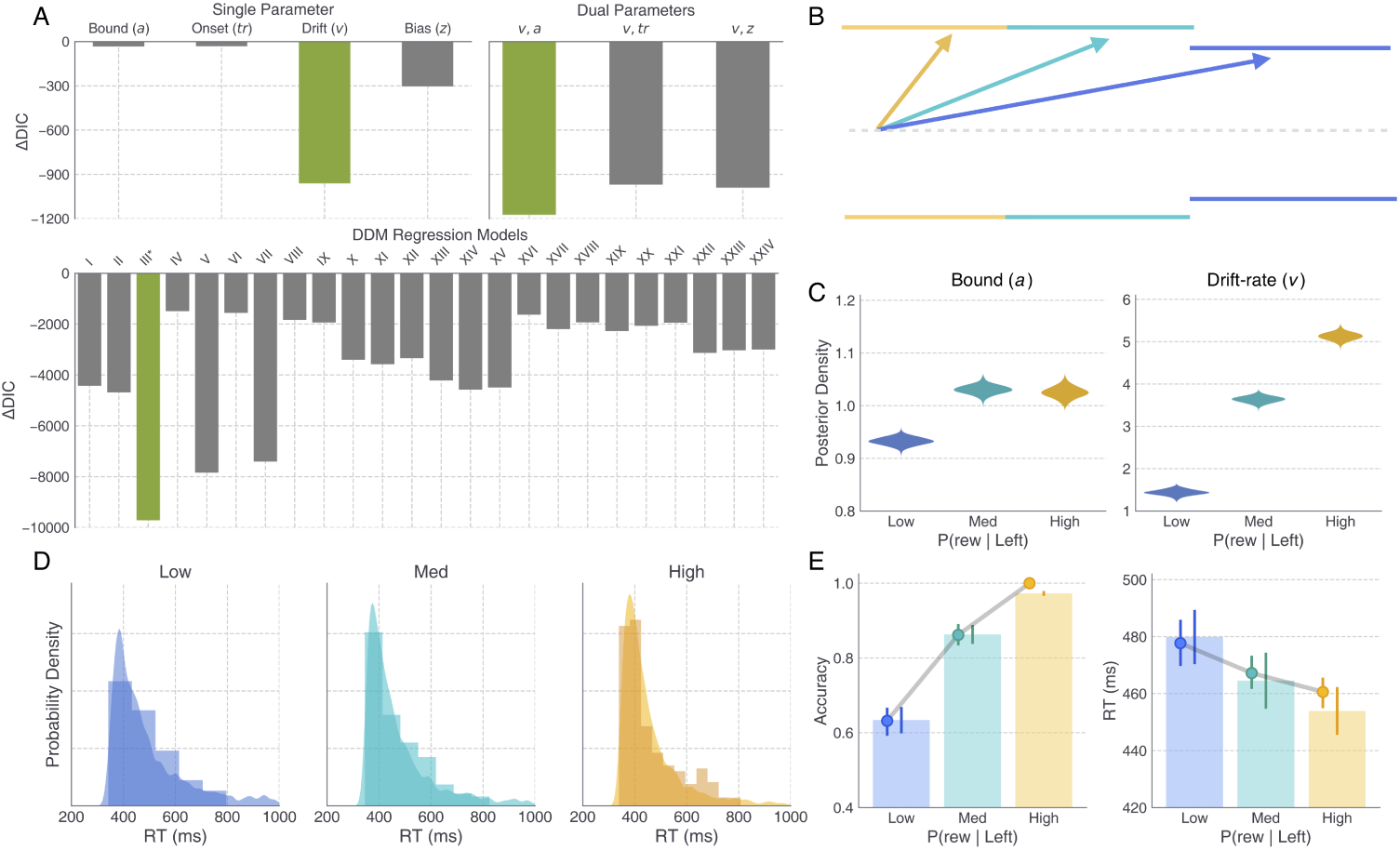
DDM fits to CBGT-simulated behavior reveals pathway-specific effects on drift rate and threshold mechanisms. A: ΔDIC scores, showing the relative goodness-of-fit of all single- and dual-parameter DDMs considered (top) and all DDM regression models considered (bottom) compared to that of the null model (all parameters held constant across conditions; see Table 2). The ΔDIC score of the best-fitting model at each stage is plotted in green. The best overall fit was provided by DDM regression model III. B: DDM schematic showing the change in *v* and *a* across low (blue), medium (cyan), and high (yellow) reward conditions, with the threshold for *L* and *R* represented as the upper and lower boundaries, respectively. C: Posterior distributions in each reward condition for *a* (eq. 1), estimated on each trial as a function of the average iMSN firing rate across left and right action channels (see *I_all_* in Figure 4C), and *v* (eq. 2), estimated on each trial as a function of the the difference between dMSN firing rates in the left and right channels (*D_L_ − D_R_* in Figure 4B). D: Histograms and kernel density estimates showing the CBGT-simulated and DDM-predicted RT distributions, respectively. E: Point plots showing the CBGT network’s average accuracy and RT across reward conditions overlaid on bars showing the DDM-predicted averages.

To investigate potential interactions between the drift rate and other parameters of the DDM, we performed another round of fits in which a second free parameter (either *a*, *tr*, or *z*), in addition to *v*, was allowed to vary across conditions (Figure 5A). Compared to alternative dual-parameter models, the combined effect of allowing *v* and *a* to vary across conditions provided the greatest improvement in model fit over the null model (ΔDIC = *−*1174.07), as well as over the best-fitting single parameter model (DIC_*v,a*_ − DIC_*v*_ = *−*213.27). While the dual *v* and *a* model significantly outperformed both alternatives (DIC_*v,a*_ − DIC_*v,t*_ = −205.89; DIC_*v,a*_ − DIC_*v,z*_ = −184.05), the second best-fitting dual parameter model, in which *v* and *z* were left free across conditions, also afforded a significant improvement over the drift-only model (DIC_*v,z*_ − DIC_*v*_ = −29.23). Thus, both *v, a* and *v, z* dual parameter models were considered in a third and final round of fits. The third round was motivated by the fact that, while behavioral fits can yield reliable and informative insights about the cognitive mechanisms engaged by a given experimental manipulation, recent studies have effectively combined behavioral observations with coincident measures of neural activity to test more precise hypotheses about the neural dynamics involved in regulating different cognitive mechanisms [34, 42, 43]. To this end, we refit the *v, a* and *v, z* models to the same simulated behavioral dataset (i.e., accuracy and RTs produced by the CBGT network) as in the previous rounds, but now with different trialwise measures of striatal activity included as regressors on one of the two free parameters in the DDM.

**Table 2.** DDM regression models and goodness-of-fit statistics. Asterisk denotes best performing model.

| | $D_L - D_R$ | $D_L - I_L$ | $I_L - I_R$ | $I_{all}$ | DIC | $\Delta DIC$ |
| --- | --- | --- | --- | --- | --- | --- |
| **I** | $v$ | $a$ | — | — | -13567.84 | -4423.64 |
| **II** | $v$ | — | $a$ | — | -13828.38 | -4684.17 |
| **\*III** | $v$ | — | — | $a$ | -18860.37 | -9716.16 |
| **IV** | — | $v$ | $a$ | — | -10636.70 | -1492.50 |
| **V** | — | $v$ | — | $a$ | -16982.35 | -7838.14 |
| **VI** | $a$ | $v$ | — | — | -10702.48 | -1558.27 |
| **VII** | — | — | $v$ | $a$ | -16547.47 | -7403.27 |
| **VIII** | $a$ | — | $v$ | — | -10979.51 | -1835.31 |
| **IX** | — | $a$ | $v$ | — | -11082.55 | -1938.34 |
| **X** | $a$ | — | — | $v$ | -12546.90 | -3402.70 |
| **XI** | — | $a$ | — | $v$ | -12719.92 | -3575.72 |
| **XII** | — | — | $a$ | $v$ | -12486.66 | -3342.46 |
| **XIII** | $v$ | $z$ | — | — | -13361.52 | -4217.32 |
| **XIV** | $v$ | — | $z$ | — | -13719.36 | -4575.16 |
| **XV** | $v$ | — | — | $z$ | -13634.12 | -4489.92 |
| **XVI** | — | $v$ | $z$ | — | -10774.88 | -1630.67 |
| **XVII** | — | $v$ | — | $z$ | -11340.47 | -2196.26 |
| **XVIII** | $z$ | $v$ | — | — | -11074.84 | -1930.64 |
| **XIX** | — | — | $v$ | $z$ | -11418.76 | -2274.56 |
| **XX** | $z$ | — | $v$ | — | -11213.79 | -2069.59 |
| **XXI** | — | $z$ | $v$ | — | -11090.96 | -1946.75 |
| **XXII** | $z$ | — | — | $v$ | -12279.57 | -3135.36 |
| **XXIII** | — | $z$ | — | $v$ | -12171.17 | -3026.96 |
| **XXIV** | — | — | $z$ | $v$ | -12144.98 | -3000.77 |

For each regression DDM (N=24 models, corresponding to 24 ways to map 2 of 4 striatal activity measures to the *v, a* and *v, z* models), one of the summary measures shown in Figure 4B-C was regressed on *v*, and another regressed on either *a* or *z*, with separate regression weights estimated for each level of reward probability. Model fit statistics are shown for each of the 24 regression models in Table 2, along with information about the neural regressors included in each model and their respective parameter dependencies. The relative goodness-of-fit afforded by all 24 regression models is visualized in Figure 5A (lower panel), identifying what we have labelled as model III as the clear winner with an overall DIC = −18860.37 and with ΔDIC = −9716.17 compared to the null model. In model III, the drift rate *v* on each action selection trial depended on the relative strength of direct pathway activation in *L* and *R* action channels (e.g., *D_L_ − D_R_*), whereas the boundary height *a* on that trial was computed as a function of the overall strength of indirect pathway activation across both channels (e.g., *I_all_*). To determine how these parameter dependencies influenced levels the boundary height and rate of evidence accumulation across levels of reward probability, the following equations were used to transform intercept and regression coefficient posteriors into posterior estimates of *a* and *v* for each condition *j*:

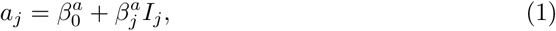

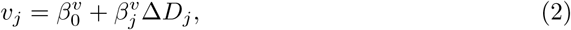

where Δ*D_j_* and *I_j_* are the mean values of *D_L_ − D_R_* and *I_all_* in condition *j* (see Figure 4B-C), 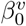 and 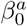 are posterior distributions for *v* and *a* intercept terms, and 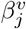 and 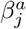 are the posterior distributions estimated for the linear weights relating *D_L_ − D_R_* and *I_all_* to *v* and *a*, respectively. The observed effects of reward probability on *v* and *a*, as mediated by trialwise changes in *D_L_ − D_R_* and *I_all_*, are schematized in Figure 5B, with conditional posteriors for each parameter plotted in Figure 5C. Consistent with best-fitting single and dual parameter models (i.e., without striatal regressors included), the weighted effect of *D_L_ − D_R_* on *v* in model III led to a significant increase in *v* across low (*µ_vLow_* = 1.43*, σ_v_Low__*= 0.06), medium (*µ_v_Med__*= 3.62*, σ_v_Med__* = 0.08), and high (*µ_v_High__* = 5.10*, σ_v_High__* = 0.09) conditions. Thus, increasing the disparity of dMSN activation between *L* and *R* action channels led to faster and more frequent leftward actions by increasing the rate of evidence accumulation towards the correct decision boundary. Also consistent with parameter estimates from the best-fitting dual parameter model (i.e., *v, a*), inclusion of trialwise values of *I_all_* led to an increase in the boundary height in the medium (*µ_a_Med__* = 1.03*, σ_a_Med__* = 0.01) and high (*µ_a_High__* = 1.02*, σ_a_High__*= 0.01) conditions compared to estimates in the low condition (*µ_a_Low__* = 0.93*, σ_a_Low__* = 0.01). However, in contrast with boundary height estimates derived from behavioral data alone, *a* estimates in model III showed no significant difference between medium and high levels of reward probability.

Examination of trialwise covariation of striatal regressors with raw choice and RT data of the network revealed additional insights into the influence of striatal dynamics on decision outcomes. Summary measures of competition between (*D_L_ − D_R_*) and within (*D_L_ − I_L_*) channels (see Figure 4B) were more predictive of choice outcome (Supplementary Figure 3A, left panels) than RT (not shown). For both *D_L_ − D_R_* (*β*=20.01, *t*(3)=30.75, *p <*0.0001) and *D_L_ − I_L_* (*β*=6.12, *t*(3)=21.37, *p <*0.0001), higher values were associated with an increased probability of choosing the higher valued action, *L*. This increased probability of choosing *L* when *D_L_ – D_R_* is high is consistent with the estimated mapping between this striatal measure and the drift rate in model III. Conversely, *I_all_* showed a stronger relationship with decision speed (Supplementary Figure 3A), with greater overall iMSN activation associated with faster response times (*β* =−2.01, *t*(3)=-53.42, *p <*0.0001). At first glance, this negative relationship between *I_all_* and RT (Supplementary Figure 3A, right) appears to contradict the role of the indirect pathway in our model as having a suppressive influence on the decision process. However, this relationship can be understood as a consequence of heightened excitatory feedback to the striatum from the thalamus on trials in which the stimulus is strong and decisions are made relatively quickly.

Next, we evaluated the extent to which the best-fitting regression model (i.e., model III) was able to account for the qualitative behavioral patterns exhibited by the CBGT network in each condition. To this end, we simulated 20,000 trials in each reward condition (each trial producing a response and RT given a parameter set sampled from the model posteriors) and compared the resulting RT distributions, along with mean speed and accuracy measures, with those produced by the CBGT model (Figure 5D,E). Parameter estimates from the best-fitting model captured the increasing positive skew of RT distributions as well as the concurrent increase in mean decision speed and accuracy with increasing reward probability.

In summary, by leveraging trialwise measures of simulated striatal MSN subpopulation dynamics to supplement RT and choice data generated by the CBGT network, we were able to 1) substantially improve the quality of DDM fits to the network’s behavior across levels of reward probability compared to models without access to neural observations and 2) identify dissociable neural signals underlying observed changes in *v* and *a* across varying levels of reward probability associated with available choices.

### 2.4 Robustness analysis

To ensure that the conclusions derived from simulations performed with the original single CBGT network were not disproportionately influenced by the chosen connectivity parameters, we performed a subsequent round of simulations and analyses aimed at assessing the robustness of our major findings. Robustness of neural and behavioral outcomes across reward conditions and of subsequent fits to the DDM were probed by re-simulating the decision experiment with multiple “subject” networks. For every projection connecting two nuclei in the CBGT network, each subject was parameterized using a randomly sampled connection efficacy (see Methods section 4.4).

Similar to the original behavioral outcomes, the sampled CBGT networks showed a trending increase in both speed and accuracy with increasing probability of reward for the better action (Figure 6A). Separate repeated-measures ANOVA tests were used to assess the statistical reliability of changes in RT and accuracy across levels of reward, revealing a marginal effect on RT, *F* (2, 28) = 3.29*, p* = 0.052, and a significant effect on accuracy, *F* (2, 28) = 465.53*, p <* 0.00001. Importantly, surrounding the subject-averaged behavioral estimates, substantial variability was observed in accuracy and RT means of individual networks. This variation in behavior across subject networks confirms that the efficacy sampling procedure had the intended effect for the purposes of assessing outcome robustness. Despite the apparent variability in behavioral means, several other patterns were conserved from the original CBGT simulations. Notably, similar to the RT distributions of the original network (see Figure 4E), the sampled network simulations led to RT distributions with an increasingly positive skew at higher levels of reward (Figure 6B), with similar ranges of RTs observed in the original and sampled networks.

**Fig 6.**
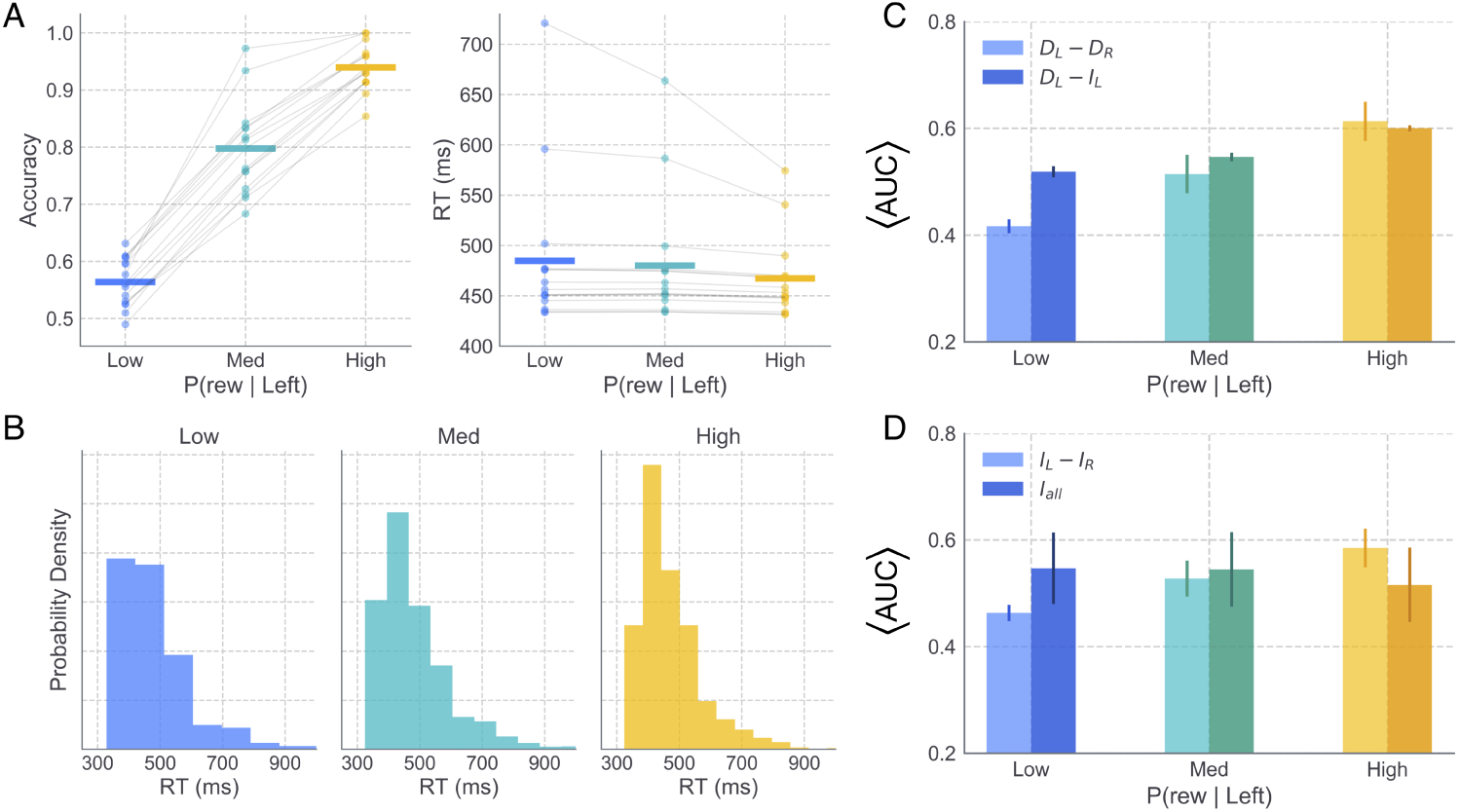
Simulated behavior and striatal influences from randomly sampled networks. A. Horizontal lines show subject-averaged accuracy (left) and correct response RT (right) means. Individual subject means are displayed as dots, connected by lines across conditions. B. Correct RT distributions in low (blue), medium (cyan), and high (yellow) reward conditions. C-D. Normalized striatal regressors, as in Figure 4C-D.

Furthermore, normalized summary measures of striatal dynamics showed consistent patterns of change across levels of reward in the sampled network simulations (Figure 6C-D) compared to those of the original network (see Figure 4B-C). As in the original simulations, the normalized difference between left and right dMSN activation (*D_L_ − D_R_*), as well as the within-channel difference between the left dMSN and iMSN activation (*D_L_ − I_L_*), both showed an increase across reward conditions (Figure 6C). Also similar to the original CBGT simulations, results with the sampled networks showed an increase in the relative activation of left and right iMSN populations (*I_L_ − I_R_*) with increasing reward, with the aggregate activation across all iMSNs (*I_all_*) staying relatively stable across conditions (Figure 6D). Consistent with the largely mirrored effects of reward on striatal summary statistics in the original and sampled networks, the time course of firing rates in striatal populations of the sampled networks showed highly similar patterns to those of the original network in each reward condition (Supplementary Figure 2).

We next sought to confirm that the mapping identified between CBGT and DDM parameters was not altered by introducing variability to the connection weights between CBGT nuclei. We found that hierarchical fits to the behavior of multiple sampled networks supported the same best-fitting single- and dual-parameter models as those selected from fits to the original single network’s behavior (Figure 7A; cf. Figure 5A). As observed in the original fits, allowing the *v* to vary across reward conditions afforded substantially better fits to the sampled network data when compared with all other single-parameter models (Figure 7A, left). Also consistent with the single network fits, allowing both the *v* and *a* to vary across reward conditions further improved the DIC score over that of the drift-only model (Figure 7A, right).

**Fig 7.**
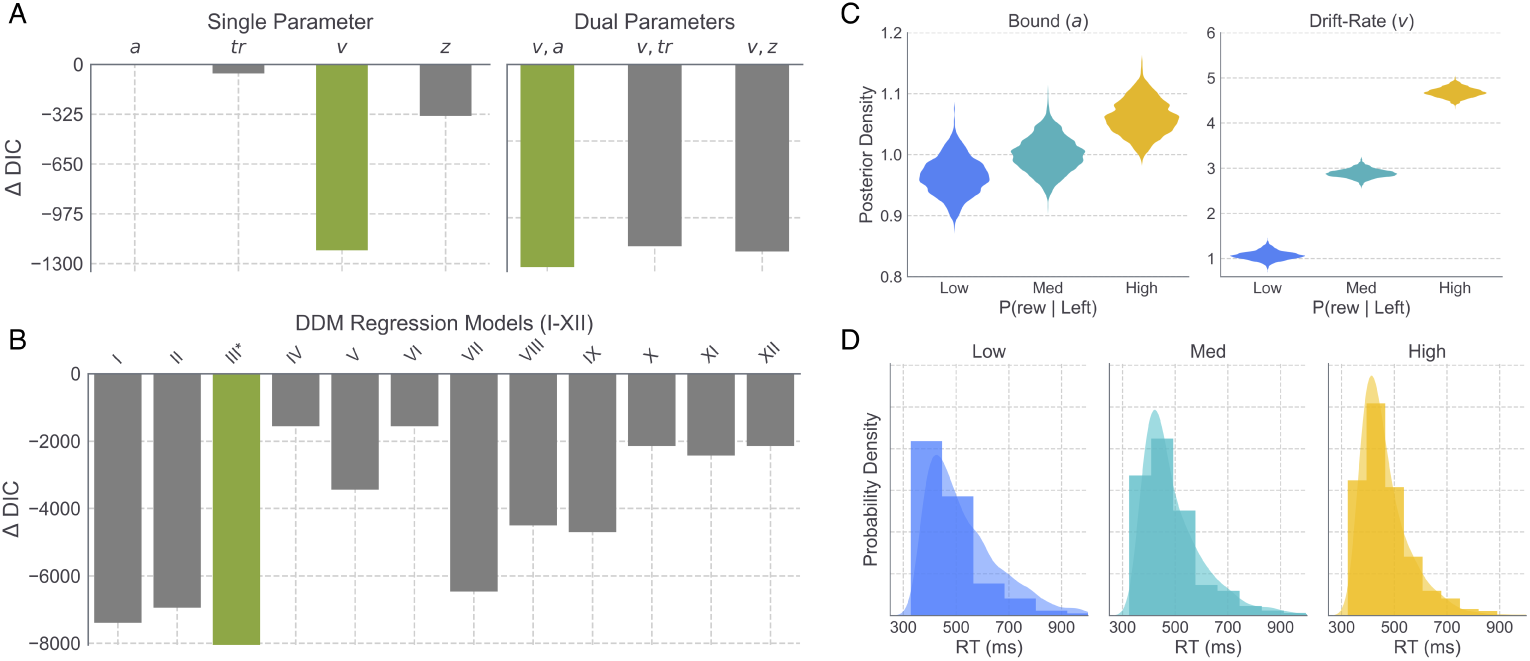
Model comparison and fits to randomly sampled network data. A. Δ*DIC* values for all single (left) and dual (right) free parameter models. B. Δ*DIC* values for DDM regression models *I − XII* associated with parameters (*v, a*). C. Posterior distributions for boundary height (*a*) and drift rate (*v*) estimated in the best-fitting regression model III. D. Correct RT distributions generated by the sampled networks (histograms) with predicted distributions from regression model III overlaid as kernel densities for each reward condition (see Figure 5 for corresponding model fit results from original network data).

Finally, we fit regression DDM models I-XII (i.e., all *v, a* models) to the data generated by the sampled networks to evaluate the robustness of the originally identified mapping between trialwise striatal dynamics and parameters of the DDM. Specifically, in the best fitting regression model for the original network (i.e., model III), trialwise fluctuations in *v* and *a* were driven by changes in *D_L_ − D_R_* and *I_all_*, respectively. Compared to alternative regression models, this same model best accounted for the behavior of the sampled networks (Figure 7B), leading to an increase in drift rate and bound with reward probability in the more-rewarded condition (Figure 7C), suggesting that this particular mapping is reasonably robust to variability in the underlying network connectivity. Similar to the original network (Supplementary Figure 3A), *D_L_ − D_R_* was highly predictive of choice outcome (*β*=29.50, *t*(3)=16.26, *p <*0.0001) in the sampled network simulations (Supplementary Figure 3B), with higher values associated with a higher probability of selecting a left action. Also consistent with the original network, *D_L_ − I_L_* was not as strongly predictive of choice outcome (*β*=4.43, *t*(3)=7.04, *p <*0.0001) as *D_L_ − D_R_*, but showed an interaction with reward level (*β*=0.71, *t*(3)=2.03, *p*=0.043; Supplementary Figure 3B). One interesting implication of this interaction between *D_L_ − I_L_* and reward level is that measures of within-channel competition between the direct and indirect pathways may be a more informative metric of an animal’s internal valuation of the corresponding action, despite sharing a weaker coupling with behavioral choice than *D_L_ − D_R_*. Indeed, given recent progress in identifying and measuring the activity of action-selective clusters of dMSN and iMSN striatal populations [44], this outcome reflects a particularly interesting testable prediction of our model. Finally, *I_all_* showed a strong negative correlation with RT in both the original and sampled network simulations (*β*=-1.38, *t*(3)=-23.46, *p <*0.0001; Supplementary Figure 3C). As noted previously, this counter-intuitive increase in speed coupled with greater overall iMSN activation can be understood as a consequence of thalamic feedback to the striatum, which becomes elevated in proportion to the motor-facilitating effects of the direct pathway on the output of the basal ganglia. As with our original simulations, parameter estimates associated with model III provided an excellent fit to RT distributions generated by the CBGT network (Figure 7D).

## 3 Discussion

Reinforcement learning in mammals alters the mapping from sensory evidence to action decisions. Here we set out to understand how this adaptive decision-making process emerges from underlying neural circuits using a modeling approach that bridges across levels of analysis, from plasticity at corticostriatal synapses to CBGT network function to quantifiable behavioral parameters [11, 12, 15, 18]. As a preliminary step, we showed that a simple, DA-mediated STDP rule alters the ratio of direct and indirect pathway corticostriatal weights within action channels in response to reward-driven feedback. With this result in hand, we simulated the network-level dynamics of CBGT circuits, as well as behavioral responses, under different levels of conflict in reward probabilities. As reward probability for the optimal target increased, the asymmetry of dMSN firing rates between action channels grew, as did the overall activity of iMSNs across both action channels. By fitting the DDM to the simulated decision behavior of the CBGT network, we found that changes in the rate of evidence accumulation tracked with the difference in dMSN population firing rates across action channels, while the the level of evidence required to trigger a decision tracked with the overall iMSN population activity. These findings show how, at least within this specific framework, plasticity at corticostriatal synapses induced by phasic changes in DA can have a multifaceted effect on cognitive decision processes.

A critical assumption of our theoretical experiments is that the CBGT pathways accumulate sensory evidence for competing actions in order to identify the most contextually appropriate response. This assumption is supported by a growing body of empirical and theoretical evidence. For example, Yartsev et al. [19] recently showed that, in rodents performing an auditory discrimination task, the anterior dorsolateral striatum satisfied three fundamental criteria for establishing causality in the evidence accumulation process: (1) inactivation of the striatum ablated the animal’s discrimination performance on the task, (2) perturbation of striatal neurons during the temporal window of evidence accumulation had predictable and reliable effects on trialwise behavioral reports, and (3) gradual ramping, proportional to the strength of evidence, was observed in both single unit and population firing rates of the striatum (however, see also [45]). Consistent with these empirical findings, Caballero et al. [16] recently proposed a novel computational framework, capturing perceptual evidence accumulation as an emergent effect of recurrent activation of competing action channels. This modeling work builds on previous studies showing how the architecture of CBGT loops is ideal for implementing a variant of the sequential probability ratio test [5, 12]. Taken together, these converging lines of evidence point to CBGT pathways as being causally involved in the accumulation of evidence for decision-making.

The idea that an accumulation of evidence algorithm can be implemented via network-level dynamics within looped circuit architectures stands in sharp contrast to cortical models of decision-making that presume a more direct isomorphism between accumulators and neural activity (for review see [20]). Early experimental work showed how population-level firing rates in area LIP displayed the same ramp-to-threshold dynamics as predicted by an evidence accumulation process [46–48]. This simple relation between algorithm and implementation has now come into question. Follow-up electrophysiological experiments showed how this population-level accumulation may, in fact, reflect the aggregation of step-functions across neurons that resemble an accumulator when summed together, yet lack accumulation properties at the level of individual units [49]. In addition, recent results from intervention studies are inconsistent with the causal role of cortical areas in the accumulation of evidence. For instance, Katz et al. [50] found that inactivation of area LIP in macaques had no effect on the ability of monkeys to discriminate the direction of motion stimuli in a standard random dot motion task. In contrast to the presumed centrality of LIP in sensory evidence accumulation, these findings and supporting reports from [51] and [52] suggest that cortical areas like LIP provide a useful proxy for the deliberation process but are unlikely to have a causal role in the decision itself.

The recent experimental [19] and theoretical [16] revelations of CBGT involvement in decision-making are particularly exciting, not only for the purposes of identifying a likely neural substrate of perceptual choice, but also for their implications for integrating accumulation-to-bound models (e.g., action selection mechanisms) with theories of RL (e.g., feedback-dependent learning of action values). We previously proposed a Believer-Skeptic framework [7] to capture the complementary roles played by the direct and indirect pathways in the feedback-dependent learning and the moment-to-moment evidence accumulation leading up to action selection (see also [22, 23]). This competition between opposing control pathways can be characterized as a debate between a Believer (direct pathway) and a Skeptic (indirect pathway), reflecting the instantaneous probability ratio of evidence in favor of executing and suppressing a given action respectively. Because the default state of the basal ganglia pathways is motor-suppressing (e.g., [53, 54]), the burden of proof falls on the Believer to present sufficient evidence for selecting a particular action. In accumulation-to-bound models like the DDM, this sequential sampling of evidence is parameterized by the drift rate. Thus, the Believer-Skeptic model assumes that this competition should be reflected, at least in part, in the rate of evidence accumulation. As for the role of learning in the Believer-Skeptic competition, multiple lines of evidence suggest that dopaminergic feedback during learning systematically biases the direct-indirect competition in a manner consistent with increasing the drift rate for more rewarding actions [7, 34, 36, 38, 55, 56]. Indeed, the STDP simulations in the current study showed opposing effects of dopaminergic feedback on corticostriatal synapses in the direct pathway when comparing across the optimal and suboptimal action channels.

In support of the biological assumptions underlying the CBGT network, several important empirical properties naturally emerged from our simulations. First, both dMSN and iMSN striatal populations were concurrently activated on each trial (see [30, 44, 57]) and exhibited gradually ramping firing rates that often saturated before the response on each trial [19, 45]. Second, in contrast with the relatively early onset of ramping activity in the striatum, recipient populations in the GPi sustained high tonic firing rates throughout most of the trial, with activity in the selected channel showing a precipitous decline near the recorded RT [27, 58, 59]. This delayed change in GPi activation is caused by the opposing influence of concurrently active dMSN and iMSN populations in each channel, such that the influence of the direct pathway on the GPi is temporarily balanced out by activation of the indirect pathway (see [27]). To represent low, medium, and high levels of reward probability conflict, we manipulated the weights of cortical input to dMSNs in each channel (see Table 4), increasing and decreasing the ratio of direct pathway weights to indirect pathway weights for *L* and *R* actions, respectively. As expected, increasing the difference in the associated reward for *L* and *R* actions led to stronger firing in *L*-dMSNs and weaker firing of *R*-dMSNs. Consistent with recently reported electrophysiological findings [30, 44], we also observed an increase in the firing of iMSNs in the *L* action channel, which in our simulations may arise from channel-specific feedback from the *L* component of the thalamus. Behaviorally, the choices of the CBGT network became both faster and more accurate (e.g., higher percentage of *L* responses) at higher levels of reward, suggesting that the observed increase in *L*-iMSN firing did not serve to delay or suppress *L* selections. These changes in neural dynamics also produced consequent changes in value-based decision behavior consistent with previous studies linking parameters of the DDM with experiential feedback.

One of the critical outcomes of the current set of experiments is the mechanistic prediction of how variation in specific neural parameters relates to changes in parameters of the DDM. Consistent with past work (see [7, 34]), the DDM fits to the CBGT-simulated behavior showed an increase in drift rate toward the higher valued decision boundary with increasing expected reward. Additionally, we found that greater disparity in the expected values of alternative actions led to an increase in the boundary height. Indeed, the co-modulation of drift rate and boundary parameters observed here has also been found in human and animal experimental studies of value-based choice [34, 36, 38]. For example, experiments with human subjects in a value-based learning task showed that selection and response speed patterns were best described by an increase in the rate of evidence for more valued targets, coupled with an upwards shift in the boundary height for all targets [36]. Moreover, in healthy human subjects, but not Parkinson’s disease patients, reward feedback was found to drive increases in both rate and boundary height parameters, effectively breaking the speed-accuracy tradeoff [36]. To identify more precise links between the relevant neural dynamics underlying the observed drift rate and boundary height effects we performed another round of model fits with striatal summary measures included as regressors to describe trial-by-trial variability. Behavioral fits were substantially improved by estimating trialwise values of drift rate as a function of the difference between *L*- and *R*-dMSN activation and trialwise values of boundary height as a function of the iMSN activation across both channels. These relationships stand both as novel predictions arising from the current study and as refinements to the competing striatal pathways framework of decision uncertainty [7, 22, 23], implying that the decision promoting component (i.e., the Believer) relies on a competition between action channels while the decision suppressing component (i.e., the Skeptic) involves a cooperative aspect across all action channels.

While our present findings provide key insights into the links between implementation mechanisms and cognitive algorithms during adaptive decision-making, they are constrained by the nature of the multi-level modeling approach itself. Our goal was to evaluate a specific hypothesis under the competing striatal pathways framework about the combined role of corticostriatal pathways in learning and decision-making, and our simulations demonstrate that strengthening corticostriatal synapses is one way that the brain can adjust striatal firing to shape the drift rate and accumulation threshold, promoting faster and more frequent selection of actions with a higher expected value. We do not presume, however, that the impacts of dopaminergic plasticity at corticostriatal synapses on striatal activity are singularly responsible for setting the drift rate during value-based decision-making. Indeed, the manipulation of a single CBGT parameter can have a compounding effect on the downstream network dynamics (e.g., amplified thalamostriatal feedback in the optimal action channel) that ultimately contributes to parametric changes at the cognitive level (e.g., an increase in the drift rate, favoring the optimal choice). Moreover, because the CBGT network has many more parameters than the DDM, many different properties of the CBGT network, aside from corticostriatial weights and measures of striatal activity, could potentially be manipulated to cause analogous behavioral patterns and inferred effects on the drift rate and boundary height parameters in the DDM. For instance, in contrast to the striatal iMSN modulation of boundary height observed in the current study, Ratcliff and Frank [18] found that simulated changes in STN firing were also capable of describing a change in the boundary height, raising the threshold in the context of high decision conflict. In fact, experimental evidence suggests the existence of both striatal [60–62] and subthalamic [42, 62, 63] mechanisms for adjusting the boundary height. It remains for future work to study how multiple mechanisms such as these work together to impact decision behavior as well as to consider more complex decision-making tasks that may help to expose distinct roles for these aspects of CBGT activity. Another open direction is to generalize our approach to include more detailed representations of neurons in CBGT populations, such as Hodgkin-Huxley-type models, and additional detail about BG neuronal subpopulations and pathways, such as distinct representations of arkypallidal and prototypical GPe neurons and the GPe projection to the striatum. Finally, while we established the robustness of our results to extensive variation of CBGT connection strengths, it is important to note that the predictions of our study depend on the assumptions and parameter choices inherent in our model construction. For instance, the STDP model used to tune corticostriatal weights treats dopamine in a simplified way, neglecting tonic dopamine as well as enhanced iMSN sensitivity to negative fluctuations in dopamine [22].

Our simulations make several novel predictions for future experiments. First, while the learning-related changes in *L* and *R* direct pathway corticostriatal weights were mirrored by the relative firing rates of *L*- and *R*-dMSNs, the iMSN firing rates expressed channel-specific differences, despite the invariance of the corticostriatal weights to iMSNs across conditions. Critically, the observed increase in iMSN firing disparity between the *L* and *R* channels in our simulations emerged due to the thalamostriatal feedback assumed in the CBGT network, where dMSN activation leads to disinhibition of the thalamus, thereby increasing excitatory feedback to both MSN subtypes within a given channel. Thus our model predicts that feedback connections from the thalamus should adhere to a channel-specific (e.g., focal) topology, which has yet to be determined by the neuroanatomical literature. In addition, if the topological predictions of our model are confirmed, the simulations presented here make the second prediction that activating the thalamic cells during the decision window should counterintuitively increase iMSN, as well as dMSN, firing rates while at the same time increasing decision speeds. One important caveat to these predictions is that thalamostriatal feedback was not incorporated into the STDP simulations used to generate the post-learning corticostriatal weights of the full CBGT network in each reward condition. Thus, additional potentiation of both dMSN and iMSN populations arising from increased thalamic gain in the optimal channel could interact with dopaminergic feedback during the learning process and thus lead to alternative downstream effects. Indeed, little is known about the role that thalamostriatal inputs play during reinforcement learning, requiring future experimental data to address these questions. Third, our simulations showed that the difference in firing rates of the dMSNs across action channels modulated the rate of value-based evidence accumulation. According to our simulations, increasing the relative magnitude of dMSN activity in the *R* channel compared to the *L* channel, for example using optogenetic stimulation of dMSNs in the dorsolateral striatum, should speed and facilitate the selection of contralateral lever presses. Choice and RT data could then be fit with the DDM to determine if the behavioral effects of laterally-biased dMSN stimulation were best described by a change in the drift rate. Analogous experiments targeting iMSNs but without channel specificity could be used similarly to evaluate our prediction that overall iMSN activity level modulates DDM boundary height.

### 3.1 Conclusion

Here we characterize the effects of dopaminergic feedback on the competition between direct and indirect CBGT pathways and how this plasticity impacts the evaluation of evidence for alternative actions during value-based choice. Using simulated neural dynamics to generate behavioral data for fitting by the DDM and determining how measures of striatal activity influence this fit, we show how the rate of evidence accumulation and the decision boundary height are modulated by the direct and indirect pathways, respectively. This multi-level modeling approach affords a unique combination of biological plausibility and mechanistic interpretability, providing a rich set of testable predictions for guiding future experimental work at multiple levels of analysis.

## 4 Methods

In our work, we generate behavioral data under several reward scenarios using a spiking *cortico-basal ganglia-thalamic (CBGT)* network, comprising neurons and synaptic connections from the key cortical and subcortical areas within the CBGT computational loops that take sensory evidence from cortex and make a decision to select one of two available responses. We fit this data using the *drift diffusion model (DDM)*, a cognitive model of decision-making that describes the accumulation-to-bound dynamics underlying the speed and accuracy of simple choice behavior [1], and we harness this fit in a hierarchical Bayesian framework (that we jointly refer to as a hierarchical DDM) to establish an upwards mapping from measures of striatal activity to DDM parameters. In this section, we briefly describe a *spike-timing dependent plasticity (STDP)* network consisting of striatal neurons and their cortical inputs, with corticostriatal synaptic plasticity driven by phasic reward signals resulting from simulated actions and their consequent dopamine release, which we use to select corticostriatal synaptic weights for our CBGT network, and then we present the details of the CBGT and DDM models along with some computational approaches that we use in simulating and analyzing them.

All simulation and analysis code reported in this work is publicly available at: https://github.com/CoAxLab/CBGT.

### 4.1 STDP network

Here we outline the general the STDP model and simulation procedures. See [29] for full details of the model implementation.

To set the corticostriatal synaptic weights across reward conditions, we consider a simplified computational model of the striatum consisting of two different populations that receive different inputs from the cortex (see Figure 1, left). Although they do not interact directly, they compete with each other to be the first to select a corresponding action. In this model, each striatal population contains two different types of units: (i) dMSNs, which facilitate action selection, and (ii) iMSNs, which suppress action selection. Each of these neurons is represented with the exponential integrate-and-fire model [64]. The inputs from the cortex to each MSN neuron within a population are generated using a collection of oscillatory Poisson processes. Each of these cortical spike trains, which we refer to as daughters, is generated from a baseline oscillatory Poisson process, which we call the mother train. We consider two different mother trains to generate the cortical daughter spike trains for the two different MSN populations. Each dMSN neuron or iMSN neuron receives input from a distinct cortical daughter train, with the corresponding transfer probabilities *p^D^* and *p^I^*, respectively. As shown in [65], the cortex to iMSN release probability exceeds that of cortex to dMSN. Hence, we set *p^D^ < p^I^*. Values of these and other parameters are selected based on properties of striatal firing determined by experimental work [66–72], on fits of network behavior to qualitative benchmarks from behavioral experiments involving action selection in a probabilistic reward environment [34], and on other computational papers involving related model components [25, 64].

These model components were coupled with an action selection criterion based on MSN spike timing, a simulated dopamine signal reflecting the comparison between actual and expected reward received upon the implementation of an action, and a learning rule for updating corticostriatal synaptic weights based on an eligibility trace determined using an STDP rule and on dopamine level. The general framework builds on previous work [22, 25, 73]. The idea of this plasticity scheme is that an eligibility trace (cf. [74]) represents a neuron’s recent spiking history and hence its eligibility to have its synapses modified, with changes in eligibility following an STDP rule that depends on both the pre- and the post-synaptic firing times. Plasticity of corticostriatal synaptic weights depends on this eligibility together with dopamine levels, which in turn depend on the reward consequences that follow neuronal spiking. One new feature is that the synaptic weight update scheme depends on the type of MSN neuron involved in the synapse, consistent with the observations that dMSNs tend to have less activity than iMSNs at resting states [66, 67, 69, 71] and are more responsive to phasic changes in dopamine than iMSNs, which are largely saturated by tonic dopamine.

Each dMSN facilitates performance of a specific action. We specify that an action occurs, and so a decision is made by the model, when at least three different dMSNs of the same population spike in a small time window of specified duration. When this condition occurs, a reward is delivered and the dopamine level is updated correspondingly based on comparison of reward level to the maximal possible action valued learned so far [75]; action values are updated via standard Q-learning. The updated dopamine level impacts all neurons in the network, depending on eligibility. Then, the spike counting and the initial window time are reset, and cortical spikes to all neurons are turned off over the next 50 *ms* before resuming again as usual. We assume that iMSN activity within a population counters the performance of the action associated with that population [76]. We implement this effect by specifying that when an iMSN in a population fires, the most recent spike fired by a dMSN in that population is suppressed. Note that this rule need not contradict observed activation of both dMSNs and iMSNs preceding a decision [31], see Subsection 2.1. For convenience, we refer to the action implemented by one population of neurons as “left” or *L* and the action selected by the other population as “right” or *R*. Based on action selection, reward delivery is probabilistic: every time that an action occurs, the reward *r_i_* is set to be 1 with probability *p_i_* or 0 otherwise, *i ∈ {L, R}*. We consider three different probabilities such that *p_L_* + *p_R_* = 1 and *p_L_ > p_R_*, keeping the action *L* as the preferred one. Specifically, we take *p_L_* = 0.85, *p_L_* = 0.75, and *p_L_* = 0.65 to allow comparison with previous results [34].

The network is integrated computationally by using the Runge-Kutta (4,5) method in Matlab (ode45) with time step *δt* = 0.01 *ms*. Different realizations lasting 15 *s* are computed to simulate variability across different subjects in a learning scenario.

### 4.2 CBGT network

The spiking CBGT network is adapted from previous work [27]. Like the STDP model described above, the CBGT network simulation is designed to decide between two actions, a left or right choice, based on incoming sensory signals (Figure 1). The full CBGT network was comprised of six interconnected brain regions (see Table 3), including populations of neurons in the cortex, striatum (STR), external segment of the globus pallidus (GPe), internal segment of the globus pallidus (GPi), subthalamic nucleus (STN), and thalamus. Because the goal of the full spiking network simulations was to probe the consequential effects of corticostriatal plasticity on the functional dynamics and emergent choice behavior of CBGT networks after learning has already occurred, CBGT simulations were conducted in the absence of any trial-to-trial plasticity, and did not include dopaminergic projections from the subtantia nigra pars compacta. Rather, corticostriatal weights were manipulated to capture the outcomes of STDP learning as simulated with the learning network (Subsection 4.1) under three different probabilistic feedback schedules (see Table 4), each maintained across all trials for that condition (N=2500 trials each).

**Table 3.**
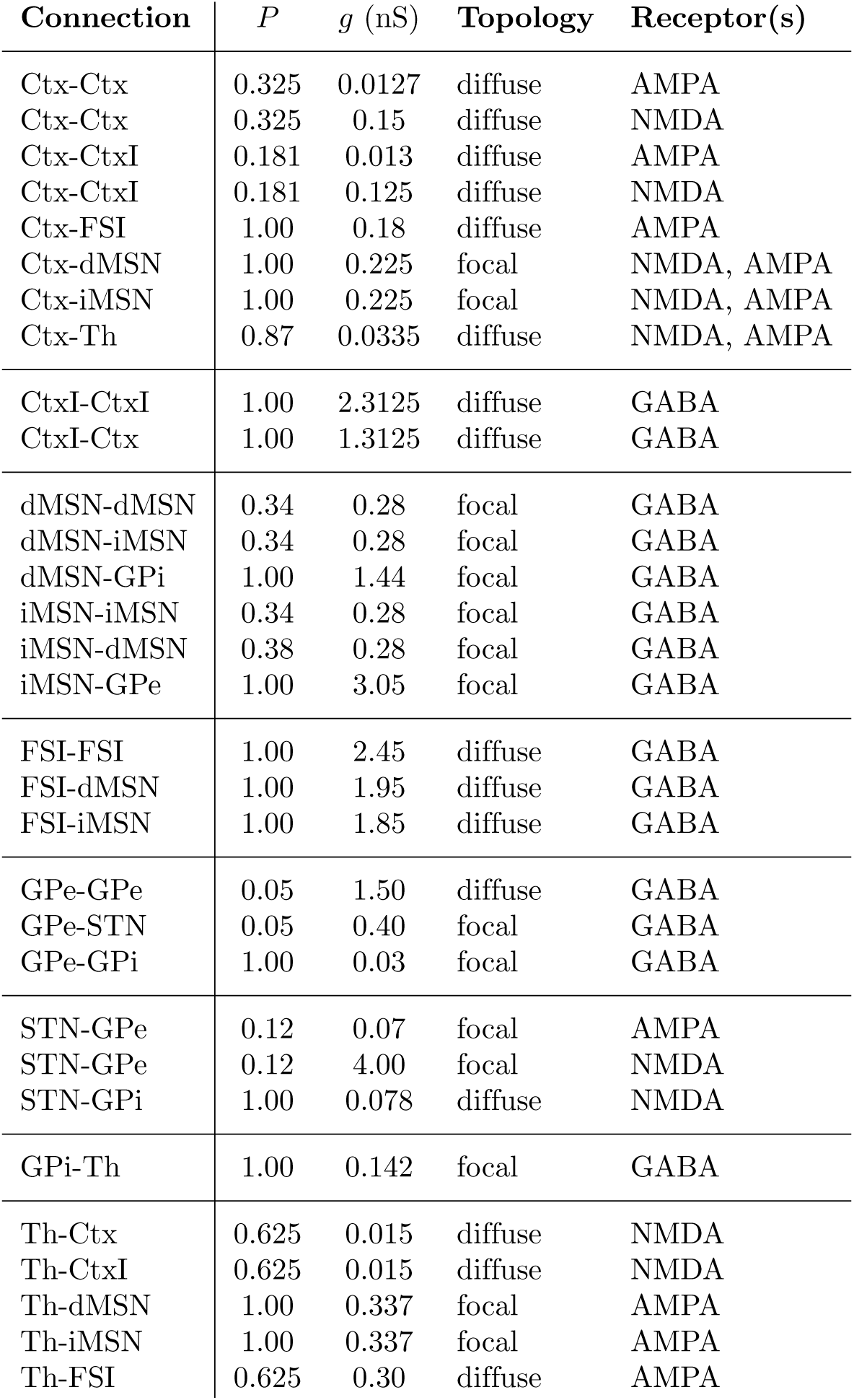
Synaptic efficacy (*g*) and probability (*P*) of connections between populations in the CBGT network, as well as postsynaptic receptor types (AMPA, NMDA, and GABA). The topology of each connection is labeled as either diffuse, to denote connections with a *P >* 0 of projecting to left and right action channels, or focal, to denote connections that were restricted to within each channel.

| Connection | $P$ | $g$ (nS) | Topology | Receptor(s) |
| --- | --- | --- | --- | --- |
| Ctx-Ctx | 0.325 | 0.0127 | diffuse | AMPA |
| Ctx-Ctx | 0.325 | 0.15 | diffuse | NMDA |
| Ctx-CtxI | 0.181 | 0.013 | diffuse | AMPA |
| Ctx-CtxI | 0.181 | 0.125 | diffuse | NMDA |
| Ctx-FSI | 1.00 | 0.18 | diffuse | AMPA |
| Ctx-dMSN | 1.00 | 0.225 | focal | NMDA, AMPA |
| Ctx-iMSN | 1.00 | 0.225 | focal | NMDA, AMPA |
| Ctx-Th | 0.87 | 0.0335 | diffuse | NMDA, AMPA |
| CtxI-CtxI | 1.00 | 2.3125 | diffuse | GABA |
| CtxI-Ctx | 1.00 | 1.3125 | diffuse | GABA |
| dMSN-dMSN | 0.34 | 0.28 | focal | GABA |
| dMSN-iMSN | 0.34 | 0.28 | focal | GABA |
| dMSN-GPi | 1.00 | 1.44 | focal | GABA |
| iMSN-iMSN | 0.34 | 0.28 | focal | GABA |
| iMSN-dMSN | 0.38 | 0.28 | focal | GABA |
| iMSN-GPe | 1.00 | 3.05 | focal | GABA |
| FSI-FSI | 1.00 | 2.45 | diffuse | GABA |
| FSI-dMSN | 1.00 | 1.95 | diffuse | GABA |
| FSI-iMSN | 1.00 | 1.85 | diffuse | GABA |
| GPe-GPe | 0.05 | 1.50 | diffuse | GABA |
| GPe-STN | 0.05 | 0.40 | focal | GABA |
| GPe-GPi | 1.00 | 0.03 | focal | GABA |
| STN-GPe | 0.12 | 0.07 | focal | AMPA |
| STN-GPe | 0.12 | 4.00 | focal | NMDA |
| STN-GPi | 1.00 | 0.078 | diffuse | NMDA |
| GPi-Th | 1.00 | 0.142 | focal | GABA |
| Th-Ctx | 0.625 | 0.015 | diffuse | NMDA |
| Th-CtxI | 0.625 | 0.015 | diffuse | NMDA |
| Th-dMSN | 1.00 | 0.337 | focal | AMPA |
| Th-iMSN | 1.00 | 0.337 | focal | AMPA |
| Th-FSI | 0.625 | 0.30 | diffuse | AMPA |

**Table 4.** Corticostriatal weights in the CBGT network across levels of reward probability. In each reward condition (rows), corresponding values of *ϕ* were used to scale the synaptic efficacy of corticostriatal inputs (*g*_Ctx-MSN_) to the direct (*D*) and indirect (*I*) pathways within the left (*L*) and right (*R*) action channels.

| P(rew\|L) | Left |  | Right |  |
| --- | --- | --- | --- | --- |
| | $\phi_L^D$ | $\phi_L^I$ | $\phi_R^D$ | $\phi_R^I$ |
| Low | 1.01 | 1.00 | 0.99 | 1.00 |
| Med. | 1.02 | 1.00 | 0.97 | 1.00 |
| High | 1.035 | 1.00 | 0.945 | 1.00 |

#### 4.2.1 Neural dynamics

To build on previous work on a two-alternative decision-making task with a similar CBGT network and to endow neurons in some BG populations with bursting capabilities, all neural units in the CBGT network were simulated using the integrate-and-fire-or-burst model [77]. Each neuron’s membrane dynamics were determined by:

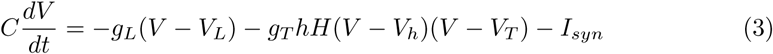

In equation 3, parameter values are *C* = 0.5 *nF*, *g_L_* = 25 *nS*, *V_L_* = *−*70 *mV*, *V_h_* = *−*0.60 *mV*, and *V_T_* = 120 *mV*. When the membrane potential reaches a boundary *V_b_*, it is reset to *V_r_*. We take *V_b_* = −50 *mV* and *V_r_* = −55 *mV*.

The middle term in the right hand side of equation 3 represents a depolarizing, low-threshold T-type calcium current that becomes available when *h* grows and when *V* is depolarized above a level *V_h_*, since *H*(*V*) is the Heaviside step function. For neurons in the cortex, striatum (both MSNs and FSIs), GPi, and thalamus, we set *g_T_* = 0, thus reducing the dynamics to the simple leaky integrate-and-fire model. For bursting units in the GPe and STN, rebound burst firing is possible, with *g_T_* set to 0.06 *nS* for both nuclei. The inactivation variable, *h*, adapts over time, decaying when *V* is depolarized and rising when *V* is hyperpolarized according to the following equations:

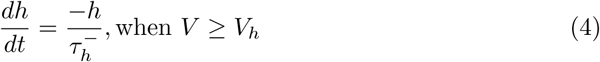

and

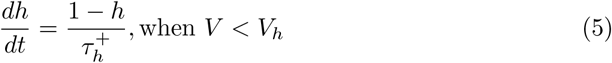

with 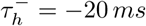 and 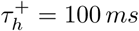 for both GPe and STN.

For all units in the model, the synaptic current *I_syn_* reflects both the synaptic inputs from other explicitly modeled populations of neurons within the CBGT network, as well as additional background inputs from sources that are not explicitly included in the model. This current is computed using the equation

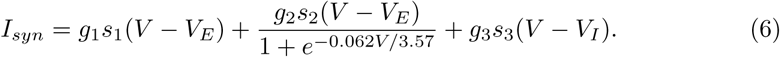

The reversal potentials are set to *V_E_* = 0 *mV* and *V_I_* = *−*70 *mV*. The synaptic current components correspond to AMPA (*g*_1_), NMDA (*g*_2_), and GABA_A_(*g*_3_) synapses. The gating variables *s_i_* for AMPA and GABA_A_ receptor-mediated currents satisfy:

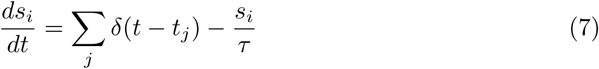

while NMDA receptor-mediated current gating obeys:

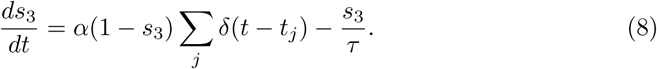

In equations 7 and 8, *t_j_* is the time of the *j^th^* spike and *α*= 0.63. The decay constant, *τ*, was 2 *ms* for AMPA, 5 *ms* for GABA A, and 100 *ms* for NMDA-mediated currents. A time delay of 0.2 *ms* was used for synaptic transmission.

#### 4.2.2 Network architecture

The CBGT network includes six of the nodes shown in Figure 1, excluding the dopaminergic projections from the substantia nigra pars compacta that are simulated in the STDP model. The membrane dynamics, projection probabilities, and synaptic weights of the network (see Table 3) were adjusted to reflect empirical knowledge about local and distal connectivity associated with different populations, as well as resting and task-related firing patterns [27, 59].

The cortex included separate populations of neurons representing sensory information for *L* (N=270) and *R* (N=270) actions that approximate the processing in the intraparietal cortex or frontal eye fields. On each trial, *L* and *R* cortical populations received excitatory inputs from an external source, sampled from a truncated normal distribution with a mean and standard deviation of 2.5 *Hz* and 0.06, respectively, with lower and upper limits of 2.4 *Hz* and 2.6 *Hz*. Critically, *L* and *R* cortical populations received the same strength of external stimulation on each trial to ensure that any observed behavioral effects across conditions were not the result of biased cortical input. Excitatory cortical neurons also formed lateral connections with other cortical neurons with a diffuse topology, or a non-zero probability of projecting to recipient neurons within and between action channels (see Table 3 for details). The cortex also included a single population of inhibitory interneurons (CtxI; N=250 total) that formed reciprocal connections with left and right sensory populations. Along with external inputs, cortical populations received diffuse ascending excitatory inputs from the thalamus (Th; N=100 per input channel).

*L* and *R* cortical populations projected to dMSN (N=100/channel) and iMSN (N=100/channel) populations in the corresponding action channel; that is, cortical signals for a *L* action projected to dMSN and iMSN cells selective for *L* actions. Both cortical populations also targeted a generic population of FSIs (N=100 total) providing widespread but asymmetric inhibition to MSNs, with stronger FSI-dMSN connections than FSI-iMSN connections [78]. Within each channel, dMSN and iMSN populations also formed recurrent and lateral inhibitory connections, with stronger inhibitory connections from iMSN to dMSN populations [78]. Striatal MSN populations also received channel-specific excitatory feedback from corresponding populations in the thalamus. Inhibitory efferent projections from the iMSNs terminated on populations of cells in the GPe, while the inhibitory efferent connections from the dMSNs projected directly to the GPi.

In addition to the descending inputs from the iMSNs, the GPe neurons (N=1000/channel, as in past work [27], reflecting divergence of connections from striatum to GPe [79, 80]) received excitatory inputs from the STN. GPe cells also formed recurrent, within channel inhibitory connections that supported stability of activity. Inhibitory efferents from the GPe terminated on corresponding populations in the the STN (i.e., long indirect pathway) and GPi (i.e., short indirect pathway). Both the projections from GPe to STN and the feedback projections from STN to GPe were sparse relative to other connection profiles in the network [81]. We did not include arkypalldal projections (i.e., feedback projections from GPe to the striatum; [82]) as it is not currently well understood how this pathway contributes to basic choice behavior.

STN populations were composed of burst-capable neurons (N=1000/channel as in GPe [27], reflecting low synaptic divergence of GPe inputs to STN [81]) with channel-specific inhibitory inputs from the GPe as well as excitatory inputs from cortex (the hyperdirect pathway). The since no cancellation signals were modeled in the experiments (see Subsection 4.2.3), the hyperdirect pathway was simplified to background input to the STN. Unlike the striatal MSNs and the GPe, the STN did not feature recurrent connections. Excitatory feedback from the STN to the GPe was assumed to be sparse but channel-specific, whereas projections from the STN to the GPi were channel-generic and caused diffuse excitation in both *L*- and *R*-encoding populations.

Populations of cells in the GPi (N=100/channel) received inputs from three primary sources: channel-specific inhibitory afferents from dMSNs in the striatum (i.e., direct pathway) and the corresponding population in the GPe (i.e., short indirect pathway), as well as excitatory projections from the STN shared across channels (i.e., long indirect and hyperdirect pathways; see Table 3). The GPi did not include recurrent feedback connections. All efferents from the GPi consisted of inhibitory projections to the motor thalamus. The efferent projections were segregated strictly into pathways for *L* and *R* actions.

Finally, *L*- and *R*-encoding populations in the thalamus were driven by two primary sources of input, integrating channel-specific inhibitory inputs from the GPi and diffuse (i.e., channel-spanning) excitatory inputs from cortex. Outputs from the thalamus delivered channel-specific excitatory feedback to corresponding dMSN and iMSN populations in the striatum as well as diffuse excitatory feedback to cortex.

#### 4.2.3 Simulations of experimental scenarios

Because the STDP simulations did not reveal strong differences in Ctx-iMSN weights across reward conditions, only Ctx-dMSN weights were manipulated across conditions in the full CBGT network simulations. In all conditions the Ctx-dMSN weights were higher in the left (higher/optimal reward probability) than in the right (lower/suboptimal reward probability) action channel (see Table 4). On each trial, external input was applied to *L*- and *R*-encoding cortical populations, each projecting to corresponding populations of dMSNs and iMSNs in the striatum, as well as to a generic population of FSIs. Critically, all MSNs also received input from the thalamus, which was reciprocally connected with cortex. Due to the suppressive effects of FSI activity on MSNs, sustained input from both cortex and thalamus was required to raise the firing rates of striatal projection neurons to levels sufficient to produce an action output. Due to the convergence of dMSN and iMSN inputs in the GPi, and their opposing influence over BG output, co-activation of these populations within a single action channel served to delay action output until activity within the direct pathway sufficiently exceeded the opposing effects of the indirect pathway [27]. The behavioral choice, as well as the time of that decision (i.e., the RT) were determined by a winner-take-all rule with the first action channel to cause the average firing rate of its thalamic population to rise above a threshold of 30 *Hz* being selected.

### 4.3 Drift Diffusion Model

To understand how altered corticostriatal weights influence decision-making behavior, we fit the simulated behavioral data from the CBGT network with a DDM [1, 83] and compared alternative models in which different parameters were allowed to vary across reward probability conditions. The DDM is an established model of simple two-alternative choice behavior, providing a parsimonious account of both the speed and accuracy of decision-making in humans and animal subjects across a wide variety of binary choice tasks [83]. It assumes that input is stochastically accumulated as the log-likelihood ratio of evidence for two alternative choices until reaching one of two decision thresholds, representing the criterion evidence for committing to a choice. Importantly, this accumulation-to-bound process affords predictions about the average accuracy, as well as the distribution of response times, under a given set of model parameters. The core parameters of the DDM include the rate of evidence accumulation, or drift rate (*v*), the distance between decision boundaries, also referred to as the threshold (*a*), the bias in the starting-point between boundaries for evidence accumulation (*z*), and a non-decision time parameter that determines when accumulation of evidence begins (*tr*), accounting for sensory and motor delays.

To narrow the subset of possible DDM models considered, DDM fits to the CBGT model behavior were conducted in three stages using a forward stepwise selection process. First, we compared models in which a single parameter in the DDM was free to vary across reward conditions. For these simulations all the DDM parameters were tested. Next, additional model fits were performed with the best-fitting model from the previous stage, but with the addition of a second free parameter. Finally, the two best fitting dual parameter models were submitted to a final round of fits in which trialwise measures of striatal activity (see Figure 4B-C) were included as regressors on the two designated parameters of the DDM. All CBGT regressors were normalized between values of 0 and 1. Each regression model included one regression coefficient capturing the linear effect of a given measure of neural activity on one of the free parameters (e.g., *a*, *v*, or *z*), as well as an intercept term for that parameter, resulting in a total of four free parameters per selected DDM parameter or 8 free parameters altogether. For example, in a model where drift rate is estimated as function of the difference between dMSN firing rates in the left and right action channels, the drift rate on trial *t* is given by 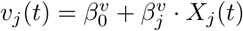, where 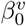 is the drift rate intercept, 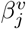 is the beta coefficient for reward condition *j*, and *X_j_*(*t*) is the observed difference in dMSN firing rates between action channels on trial *t* in condition *j*. A total of 24 separate regression models were fit, testing all possible combinations between the two best-fitting dual parameter models and the four measures of striatal activity summarized in Figure 4B-C.

Fits of the DDM were performed using HDDM (see [84] for details), an open source Python package for Bayesian estimation of DDM parameters. Each model was fit by drawing 2000 Markov Chain Monte-Carlo (MCMC) samples from the joint posterior probability distribution over all parameters, with acceptance based on the likelihood (see [85]) of the observed accuracy and RT data given each parameter set. A burn-in period of 1200 samples was implemented to ensure that model selection was not influenced by samples drawn prior to convergence. Sampling chains were also visually inspected for signs of convergence failure; however, parameters in all models showed normally distributed posterior distributions with little autocorrelation between samples suggesting that sampling parameters were sufficient for convergence. The prior distributions used to initialize all DDM parameters included in the fits can be found in [84].

### 4.4 Network sampling procedure

To assess the robustness of our findings to alterations in the connection weights used in the CBGT network, we repeated the simulation and fitting procedures described in sections 4.2.3 and 4.3, respectively, with additional variability in the parameterization of CBGT connection strengths. The functional outputs of the spiking CBGT network depend on two critical parameters:

1. *P_ij_*: the probability that a cell in population *i* projects to a cell in population *j*,
2. *g_ij_*: the conductance strength associated with inputs from cells in population *i* to cells in population *j*.

The product of these two values can be taken to represent the weight (*ω_ij_*) of connections from population *i* to population *j*. The core simulations in this work were performed using a single *ω_ij_* for each connection that was held constant for all simulated trials. We also introduced variability in *ω_ij_* in additional simulations to explore robustness of results to variability in the underlying network connectivity.

For our robustness check, rather than simulating 2500 trials per condition with a single set of CBGT connectivity parameters, we generated multiple “subject” networks, simulating 200 trials per condition with each. For each subject *k* (*N* = 15), *g*(*k*)_*ij*_ was independently sampled from a normal distribution with mean 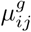 and standard deviation 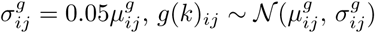. For each connection included in the random sampling procedure, *µ_ij_* was set equal to the value of *g_ij_* in Table 3. The total strength of the connection from population *i* to population *j* in subject network *k* was given by the equation *ω*(*k*)_*ij*_ = *P_ij_ ⋅ g*(*k*)_*ij*_.

As in the original single network simulations, the dopaminergic effects on synaptic strengths of corticostriatal inputs to left (*L*, optimal) and right (*R*, suboptimal) action channels were determined by scaling the baseline value of *g*_Ctx-MSN_ by the proportional change in synaptic strength *φ* observed in the STDP simulations in reward condition *C* (see Table 4). Thus, for subject network *k*, the weight of Ctx − dMSN connections in the left 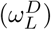 and right 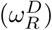 action channels were given by the following equations:

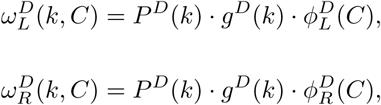

where *P ^D^* is the connection probability of cortical inputs to L and R dMSN populations (see Table 3), *g^D^*(*k*) is sampled synaptic efficacy for subject network *k*, and *φ*(*C*) is the proportional change due to DA-mediated STDP in condition *C* indicated in Table 4. Because Ctx − iMSN weights were less sensitive to dopaminergic feedback in STDP simulations, manipulations of corticostriatal weights across levels of reward were restricted to Ctx *−* dMSN connections.

### 4.5 Hierarchical fits to sampled networks

To confirm that variability in the underlying connection weights of the network did not alter the upwards mapping between CBGT and DDM parameters, all single and dual static parameter models and regression models I-XII were re-fit to the data generated by the sampled networks. In contrast to the original model fits, which were performed on data from a single network, model fits to the sampled network data were performed hierarchically. Hierarchical model fits allow for simultaneous and conditional estimation of parameters at the individual and group levels. At the individual level, samples were drawn from the joint posterior distribution given each individual subject’s data. At the group level, these individual subject samples were aggregated to derive estimates of the mean and variance for each parameter. Finally, these estimates of group-level summary statistics served as prior distributions at the subject-level, reducing the weight of outlier subjects that deviate significantly from the group.

All single and dual parameter models were defined in the same way as those fit to the original network, but with all free parameters estimated for each individual network and with corresponding estimates of the mean and variance of each parameter at the group level. Regression DDM fits were also performed hierarchically, with a random intercept for drift rate and boundary-height estimated for each individual subject network, and with regression weights estimated as fixed effects (i.e., shared across subject networks at the group level).

## Acknowledgments

CV is supported by the Ministerio de Economía, Industria y Competitividad (MINECO), the Agencia Estatal de Investigación (AEI), and the European Regional Development Funds (ERDF) through projects MTM2014-54275-P, MTM2015-71509-C2-2-R and MTM2017-83568-P (AE/ERDF,EU). JR received support from NSF awards DMS 1516288, 1612913 (CRCNS), and 1724240 (CRCNS). TV received support from NSF CAREER award 1351748. The research was sponsored in part by the U.S. Army Research Laboratory, including work under Cooperative Agreement Number W911NF-10-2-0022, and the views espoused are not official policies of the U.S. Government.

## Supplementary Information

**Supp. Figure 1.**
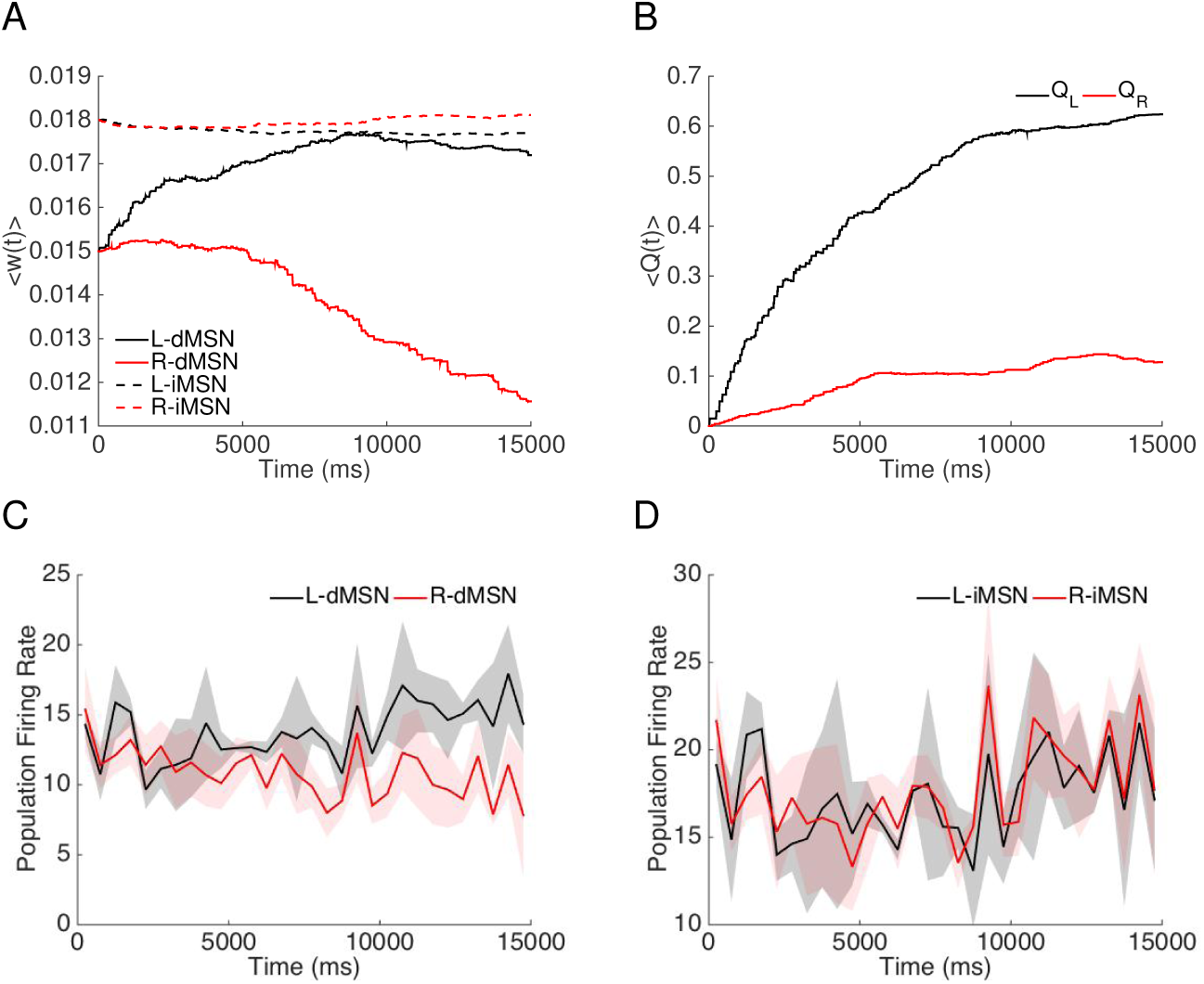
Time courses of corticostriatal synapse weights and firing rates when the rewards are constant in time (*r_L_*(*t*) = 0.7 and *r_R_*(*t*) = 0.1). A: Averaged weights over 7 different realizations and over each of the four specific populations of neurons, which are dMSN selecting action *L* (solid black); dMSN selecting action *R* (solid red); iMSN countering action *L* (dashed black); iMSN countering action *R* (dashed red). B: Averaged evolution of the action values *Q_L_* (black trace) and *Q_R_* (red trace) over 7 different realizations. C: Average neuronal firing rate (spike count within a time bin divided by bin duration) across neurons in the dMSN population selecting action *L* (black) and *R* (red), respectively, over time. D: Firing rates of the iMSN populations countering actions *L* (black) and *R* (red) over time. Data in C,D was discretized into 50 *ms* bins. The transparent regions depict standard deviations.

**Supp. Figure 2.**
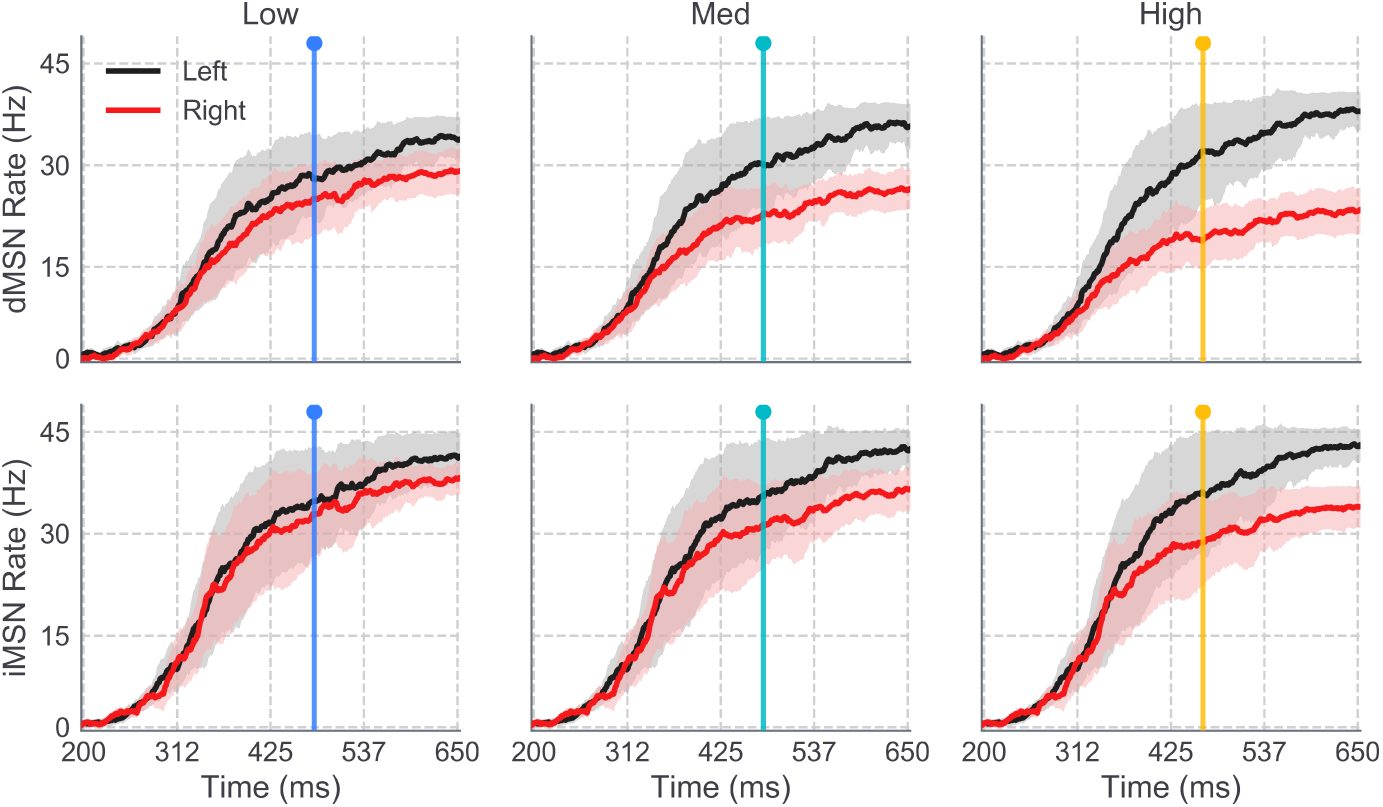
Population firing rates of sampled network MSNs. Timecourse of average firing rates of dMSN (upper) and iMSN (lower) populations in left (black) and right (red) action channels are shown for low (left), medium (middle), and high (right) reward conditions.

**Supp. Figure 3.**
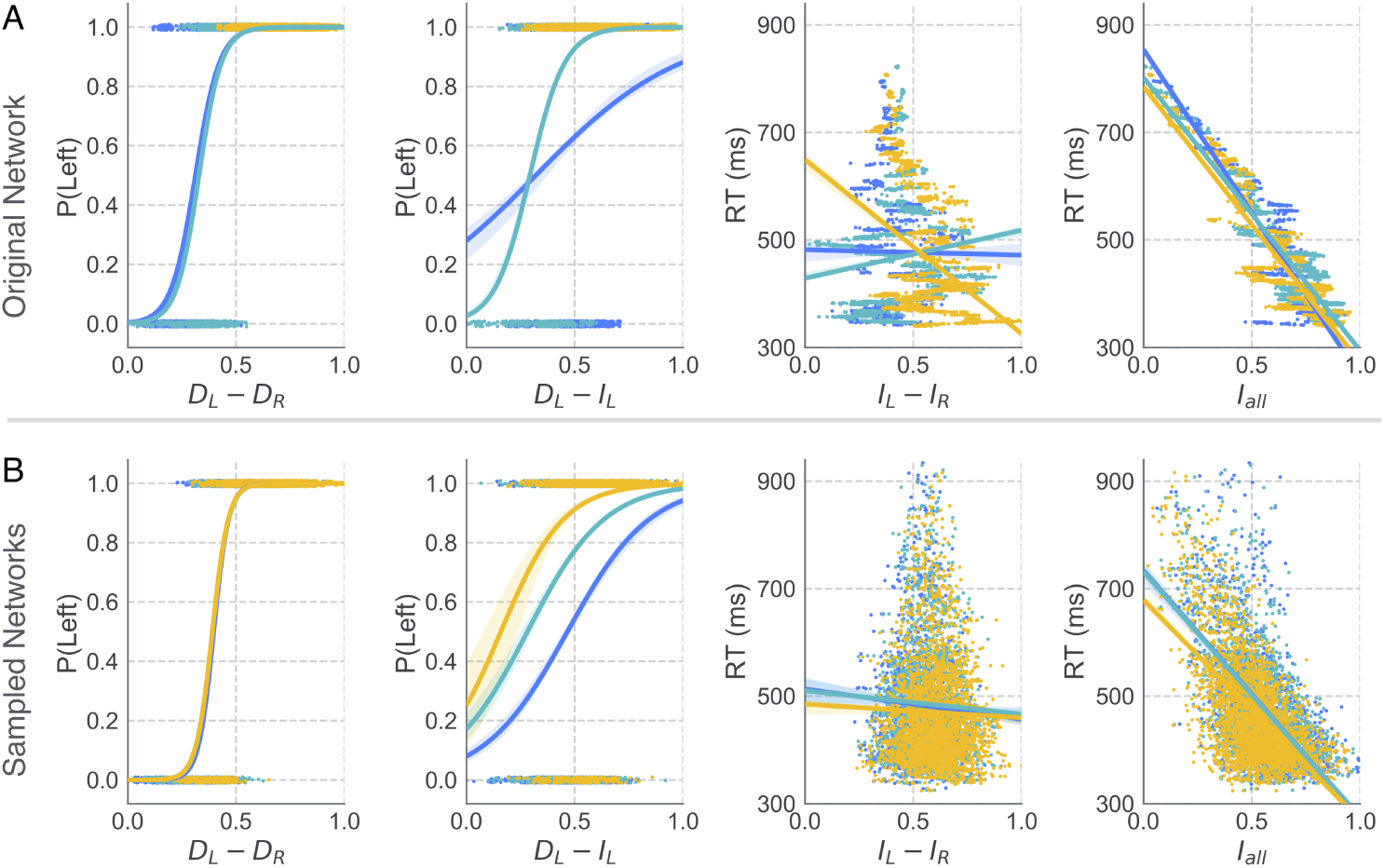
Effects of striatal dynamics on behavior. A. Simulated decision outcomes (left, center-left) and RTs (right, center-right) from the original CBGT network are plotted as a function of the four striatal summary statistics used as predictors of trialwise changes in DDM parameters. Choice outcome is plotted as a function of direct pathway measures, *D_L_ − D_R_* (left) and *D_L_ − I_L_* (center-left); RT as a function of *I_L_ − I_R_* (center-right) and *I_all_* (right). Each dot represents data from an individual trial in the low (blue), medium (cyan), and high (yellow) reward conditions. Logistic (left, center-left) and linear (right, center-right) regression predictions for each condition are shown as lines. B. The same data as shown in panel A from multiple (N=15) randomly sampled CBGT networks.

## References

1. Ratcliff R. A theory of Memory Retrival. Psychol Rev. 1978;85(2):59–108.

2. Sutton RS, Barto AG, Book aB. Reinforcement Learning : An Introduction. 1998;.

3. Rescorla RA, Wagner AR, et al. A theory of Pavlovian conditioning: Variations in the effectiveness of reinforcement and nonreinforcement. Classical conditioning II: Current research and theory. 1972;2:64–99.

4. Doya K. Modulators of decision making. Nat Neurosci. 2008;11(4):410–416.

5. Bogacz R, Gurney K. The basal ganglia and cortex implement optimal decision making between alternative actions. Neural computation. 2007;19(2):442–477.

6. Balleine BW, Delgado MR, Hikosaka O. The role of the dorsal striatum in reward and decision-making. J Neurosci. 2007;27(31):8161–8165.

7. Dunovan K, Verstynen T. Believer-Skeptic meets Actor-Critic: Rethinking the role of basal ganglia pathways during decision-making and reinforcement learning. Frontiers in neuroscience. 2016;10:106.

8. Nonomura S, Nishizawa K, Sakai Y, Kawaguchi Y, Kato S, Uchigashima M, et al. Monitoring and Updating of Action Selection for Goal-Directed Behavior through the Striatal Direct and Indirect Pathways. Neuron. 2018;99(6):1302–1314.e5.

9. Marr D, Poggio T. From understanding computation to understanding neural circuitry. 1976;.

10. Krakauer JW, Ghazanfar AA, Gomez-Marin A, MacIver MA, Poeppel D. Neuroscience needs behavior: correcting a reductionist bias. Neuron. 2017;93(3):480–490.

11. Simen P, Cohen JD, Holmes P. Rapid decision threshold modulation by reward rate in a neural network. Neural Netw. 2006;19(8):1013–1026.

12. Bogacz R. Optimal decision-making theories: linking neurobiology with behaviour. Trends in cognitive sciences. 2007;11(3):118–125.

13. Draglia V, Tartakovsky AG, Veeravalli VV. Multihypothesis sequential probability ratio tests. I. Asymptotic optimality. IEEE Transactions on Information Theory. 1999;45(7):2448–2461.

14. Baum CW, Veeravalli VV. A sequential procedure for multihypothesis testing. IEEE Transactions on Information Theory. 1994;40(6).

15. Bogacz R, Larsen T. Integration of reinforcement learning and optimal decision-making theories of the basal ganglia. Neural computation. 2011;23(4):817–851.

16. Caballero JA, Humphries MD, Gurney KN. A probabilistic, distributed, recursive mechanism for decision-making in the brain. PLoS Comput Biol. 2018;14(4):e1006033.

17. Frank MJ. Linking Across Levels of Computation in Model-Based Cognitive Neuroscience. In: An Introduction to Model-Based Cognitive Neuroscience. Springer, New York, NY; 2015. p. 159–177.

18. Ratcliff R, Frank MJ. Reinforcement-Based Decision Making in Corticostriatal Circuits: Mutual Constraints by Neurocomputational and Diffusion Models. Neural Comput. 2012;24:1186–1229.

19. Yartsev MM, Hanks TD, Yoon AM, Brody CD. Causal contribution and dynamical encoding in the striatum during evidence accumulation. Elife. 2018;7:e34929.

20. Gold JI, Shadlen MN. The neural basis of decision making. Annu Rev Neurosci. 2007;30(30):535–561.

21. Dunovan K, Lynch B, Molesworth T, Verstynen T. Competing basal ganglia pathways determine the difference between stopping and deciding not to go. Elife. 2015;4:e08723.

22. Mikhael JG, Bogacz R. Learning Reward Uncertainty in the Basal Ganglia. PLoS Comput Biol. 2016;12(9):e1005062.

23. Bariselli S, Fobbs W, Creed M, Kravitz A. A competitive model for striatal action selection. Brain research. 2018;.

24. Gurney KN, Humphries MD, Redgrave P. A New Framework for Cortico-Striatal Plasticity: Behavioural Theory Meets In Vitro Data at the Reinforcement-Action Interface. PLOS Biology. 2015;13(1):1–25. doi:10.1371/journal.pbio.1002034.

25. Baladron J, Nambu A, Hamker FH. The subthalamic nucleus-external globus pallidus loop biases exploratory decisions towards known alternatives: a neuro-computational study. European Journal of Neuroscience. 2017; p. 1–14. doi:10.1111/ejn.13666.

26. Schultz W, Apicella P, Scarnati E, Ljungberg T. Neuronal activity in monkey ventral striatum related to the expectation of reward. J Neurosci. 1992;12(12):4595–4610.

27. Wei W, Rubin JE, Wang XJ. Role of the indirect pathway of the basal ganglia in perceptual decision making. J Neurosci. 2015;35(9):4052–4064.

28. Wiecki TV, Frank MJ. A computational model of inhibitory control in frontal cortex and basal ganglia. Psychol Rev. 2013;120(2):329–355.

29. Vich C, Dunovan K, Verstynen T, Rubin J. Corticostriatal synaptic weight evolution in a two-alternative forced choice task. bioRxiv. 2019;.

30. Donahue CH, Liu M, Kreitzer A. Distinct value encoding in striatal direct and indirect pathways during adaptive learning. bioRxiv. 2018;doi:10.1101/277855.

31. Cui G, Jun SB, Jin X, Pham MD, Vogel SS, Lovinger DM, et al. Concurrent activation of striatal direct and indirect pathways during action initiation. Nature. 2013;494(7436):238–242.

32. Tecuapetla F, Matias S, Dugue GP, Mainen ZF, Costa RM. Balanced activity in basal ganglia projection pathways is critical for contraversive movements. Nature communications. 2014;5:4315.

33. Tecuapetla F, Jin X, Lima SQ, Costa RM. Complementary contributions of striatal projection pathways to action initiation and execution. Cell. 2016;166(3):703–715.

34. Frank MJ, Gagne C, Nyhus E, Masters S, Wiecki TV, Cavanagh JF, et al. fMRI and EEG predictors of dynamic decision parameters during human reinforcement learning. J Neurosci. 2015;35(2):485–494.

35. Parker JG, Marshall JD, Ahanonu B, Wu YW, Kim TH, Grewe BF, et al. Diametric neural ensemble dynamics in parkinsonian and dyskinetic states. Nature. 2018;557(7704):177.

36. Manohar SG, Chong TTJ, Apps MAJ, Batla A, Stamelou M, Jarman PR, et al. Reward Pays the Cost of Noise Reduction in Motor and Cognitive Control. Curr Biol. 2015;25(13):1707–1716.

37. Polania R, Krajbich I, Grueschow M, Ruff CC. Neural Oscillations and Synchronization Differentially Support Evidence Accumulation in Perceptual and Value-Based Decision Making. Neuron. 2014;82(3):709–720.

38. Afacan-Seref K, Steinemann NA, Blangero A, Kelly SP. Dynamic Interplay of Value and Sensory Information in High-Speed Decision Making. Current Biology. 2018;28(5):795–802.

39. Gardner MPH, Conroy JS, Shaham MH, Styer CV, Schoenbaum G. Lateral Orbitofrontal Inactivation Dissociates Devaluation-Sensitive Behavior and Economic Choice. Neuron. 2017;96(5):1192–1203.e4.

40. Jahfari S, Ridderinkhof KR, Collins AGE, Knapen T, Waldorp L, Frank MJ. Cross-task contributions of fronto-basal ganglia circuitry in response inhibition and conflict-induced slowing. bioRxiv. 2017; p. 199299.

41. Burnham KP, Anderson DR. Model Selection and Inference: A Practical Information-Theoretic Approach. vol. 80; 1998.

42. Herz DM, Zavala BA, Bogacz R, Brown P. Neural correlates of decision thresholds in the human subthalamic nucleus. Current Biology. 2016;26(7):916–920.

43. Herz DM, Little S, Pedrosa DJ, Tinkhauser G, Cheeran B, Foltynie T, et al. Mechanisms Underlying Decision-Making as Revealed by Deep-Brain Stimulation in Patients with Parkinson’s Disease. Current Biology. 2018;28(8):1169–1178.

44. Klaus A, Martins GJ, Paixao VB, Zhou P, Paninski L, Costa RM. The Spatiotemporal Organization of the Striatum Encodes Action Space. Neuron. 2017;95(5):1171–1180.e7. doi:https://doi.org/10.1016/j.neuron.2017.08.015.

45. Ding L, Gold JI. Caudate encodes multiple computations for perceptual decisions. J Neurosci. 2010;30(47):15747–15759.

46. Shadlen MN, Newsome WT. Neural basis of a perceptual decision in the parietal cortex (area LIP) of the rhesus monkey. Journal of neurophysiology. 2001;86(4):1916–1936.

47. Kiani R, Shadlen MN. Representation of confidence associated with a decision by neurons in the parietal cortex. science. 2009;324(5928):759–764.

48. Churchland AK, Kiani R, Shadlen MN. Decision-making with multiple alternatives. Nat Neurosci. 2008;11(6):693–702.

49. Latimer KW, Yates JL, Meister MLR, Huk AC, Pillow JW. Single-trial spike trains in parietal cortex reveal discrete steps during decision-making. Science. 2015;349(6244):184–187.

50. Katz LN, Yates JL, Pillow JW, Huk AC. Dissociated functional significance of decision-related activity in the primate dorsal stream. Nature. 2016;535(7611):285–288.

51. Licata AM, Kaufman MT, Raposo D, Ryan MB, Sheppard JP, Churchland AK. Posterior Parietal Cortex Guides Visual Decisions in Rats. J Neurosci. 2017;37(19):4954–4966.

52. Erlich JC, Brunton BW, Duan CA, Hanks TD, Brody CD. Distinct effects of prefrontal and parietal cortex inactivations on an accumulation of evidence task in the rat. Elife. 2015;4:e05457.

53. Alexander GE, Crutcher MD. Functional architecture of basal ganglia circuits: neural substrates of parallel processing. Trends Neurosci. 1990;13(7):266–271.

54. Wichmann T, DeLong MR. Functional and pathophysiological models of the basal ganglia. Current Opinion in Neurobiology. 1996;6(6):751–758. doi:https://doi.org/10.1016/S0959-4388(96)80024-9.

55. Pedersen ML, Frank MJ, Biele G. The drift diffusion model as the choice rule in reinforcement learning. Psychonomic bulletin & review. 2017;24(4):1234–1251.

56. Collins AGE, Frank MJ. Opponent actor learning (OpAL): modeling interactive effects of striatal dopamine on reinforcement learning and choice incentive. Psychol Rev. 2014;121(3):337–366.

57. Cui Y, Paillie V, Xu H, Genet S, Delord B, Fino E, et al. Endocannabinoids mediate bidirectional striatal spike-timing-dependent plasticity. Journal of Physiology. 2015;593(13):2833–2849. doi:10.1113/JP270324.

58. Schmidt R, Leventhal DK, Mallet N, Chen F, Berke JD. Canceling actions involves a race between basal ganglia pathways. Nat Neurosci. 2013;16(8):1118–1124.

59. Lo CC, Wang XJ. Cortico-basal ganglia circuit mechanism for a decision threshold in reaction time tasks. Nat Neurosci. 2006;9(7):956–963.

60. Forstmann BU, Dutilh G, Brown S, Neumann J, von Cramon DY, Ridderinkhof KR, et al. Striatum and pre-SMA facilitate decision-making under time pressure. Proc Natl Acad Sci U S A. 2008;105(45):17538–17542.

61. Forstmann BU, Anwander A, Schäfer A, Neumann J, Brown S, Wagenmakers EJ, et al. Cortico-striatal connections predict control over speed and accuracy in perceptual decision making. Proc Natl Acad Sci U S A. 2010;107(36):15916–15920.

62. Bogacz R, Wagenmakers EJ, Forstmann BU, Nieuwenhuis S. The neural basis of the speed-accuracy tradeoff. Trends in neurosciences. 2010;33(1):10–16.

63. Herz DM, Tan H, Brittain JS, Fischer P, Cheeran B, Green AL, et al. Distinct mechanisms mediate speed-accuracy adjustments in cortico-subthalamic networks. Elife. 2017;6.

64. Fourcaud-Trocm_e N, Hansel D, van Vreeswijk C, Brunel N. How spike generation mechanisms determine the neuronal response to fluctuating inputs. J Neurosci. 2003;23(37):11628–11640.

65. Kreitzer AC, Malenka RC. Striatal Plasticity and Basal Ganglia Circuit Function. Neuron. 2008;60(4):543–554. doi:https://doi.org/10.1016/j.neuron.2008.11.005.

66. Richfield EK, Penney JB, Young AB. Anatomical and affinity state comparisons between dopamine D1 and D2 receptors in the rat central nervous system. Neuroscience. 1989;30(3):767–777. doi:https://doi.org/10.1016/0306-4522(89)90168-1.

67. Gonon F. Prolonged and Extrasynaptic Excitatory Action of Dopamine Mediated by D1 Receptors in the Rat Striatum In Vivo. Journal of Neuroscience. 1997;17(15):5972–5978.

68. Mallet N, Ballion B, Le Moine C, Gonon F. Cortical Inputs and GABA Interneurons Imbalance Projection Neurons in the Striatum of Parkinsonian Rats. Journal of Neuroscience. 2006;26(14):3875–3884. doi:10.1523/JNEUROSCI.4439-05.2006.

69. Dreyer JK, Herrik KF, Berg RW, Hounsgaard JD. Influence of Phasic and Tonic Dopamine Release on Receptor Activation. Journal of Neuroscience. 2010;30(42):14273–14283. doi:10.1523/JNEUROSCI.1894-10.2010.

70. Flores-Barrera E, Vizcarra-Chacón B, Tapia D, Bargas J, Galarraga E. Different corticostriatal integration in spiny projection neurons from direct and indirect pathways. Frontiers in Systems Neuroscience. 2010;4:15. doi:10.3389/fnsys.2010.00015.

71. Keeler J, Pretsell D, Robbins T. Functional implications of dopamine D1 vs. D2 receptors: a ‘prepare and select’model of the striatal direct vs. indirect pathways. Neuroscience. 2014;282:156–175.

72. Escande MV, Taravini IRE, Zold CL, Belforte JE, Murer MG. Loss of Homeostasis in the Direct Pathway in a Mouse Model of Asymptomatic Parkinson’s Disease. Journal of Neuroscience. 2016;36(21):5686–5698. doi:10.1523/JNEUROSCI.0492-15.2016.

73. Izhikevich EM. Dynamical systems in neuroscience: the geometry of excitability and bursting. Computational Neuroscience. Cambridge, MA: MIT Press; 2007.

74. Shindou T, Shindou M, Watanabe S, Wickens J. A silent eligibility trace enables dopamine-dependent synaptic plasticity for reinforcement learning in the mouse striatum. Eur J Neurosci. 2018;.

75. Roesch MR, Calu DJ, Schoenbaum G. Dopamine neurons encode the better option in rats deciding between differently delayed or sized rewards. Nature Neuroscience. 2007;10:1615–1624.

76. Roseberry TK, Lee AM, Lalive AL, Wilbrecht L, Bonci A, Kreitzer AC. Cell-Type-Specific Control of Brainstem Locomotor Circuits by Basal Ganglia. Cell. 2016;164(3):526–537. doi:https://doi.org/10.1016/j.cell.2015.12.037.

77. Smith GD, Cox CL, Sherman SM, Rinzel J. Fourier analysis of sinusoidally driven thalamocortical relay neurons and a minimal integrate-and-fire-or-burst model. J Neurophysiol. 2000;83(1):588–610.

78. Gittis AH, Nelson AB, Thwin MT, Palop JJ, Kreitzer AC. Distinct roles of GABAergic interneurons in the regulation of striatal output pathways. J Neurosci. 2010;30(6):2223–2234.

79. Fox C, Rafols J. The striatal efferents in the globus pallidus and in the substantia nigra. Research Publications-Association for Research in Nervous and Mental Disease. 1976;55:37–55.

80. Smith Y, Bevan M, Shink E, Bolam J. Microcircuitry of the direct and indirect pathways of the basal ganglia. Neuroscience. 1998;86(2):353–387.

81. Steiner LA, Tomás FJB, Planert H, Alle H, Vida I, Geiger JR. Connectivity and dynamics underlying synaptic control of the Subthalamic Nucleus. Journal of Neuroscience. 2019; p. 1642–18.

82. Mallet N, Micklem BR, Henny P, Brown MT, Williams C, Bolam JP, et al. Dichotomous Organization of the External Globus Pallidus. Neuron. 2012;74(6):1075–1086.

83. Ratcliff R, Smith PL, Brown SD, McKoon G. Diffusion Decision Model: Current Issues and History. Trends Cogn Sci. 2016;20(4):260–281.

84. Wiecki TV, Sofer I, Frank MJ. HDDM: hierarchical bayesian estimation of the drift-diffusion model in python. Frontiers in neuroinformatics. 2013;7:14.

85. Navarro DJ, Fuss IG. Fast and accurate calculations for first-passage times in Wiener diffusion models. J Math Psychol. 2009;53(4):222–230.

